# ENDOTHELIAL PROX1 INDUCES BLOOD-BRAIN BARRIER DISRUPTION IN THE CENTRAL NERVOUS SYSTEM

**DOI:** 10.1101/2024.10.03.616513

**Authors:** Sara González-Hernández, Ryo Sato, Yuya Sato, Chang Liu, Wenling Li, Chengyu Liu, Sadhana Jackson, Yoshiaki Kubota, Yoh-suke Mukouyama

**Author notes:** Author for correspondence: Yoh-suke Mukouyama. Conflict-of-interest statement: The authors have declare that no conflict of interest exists.

## Abstract

The central nervous system (CNS) parenchyma has conventionally been believed to lack lymphatic vasculature, likely due to a non-permissive microenvironment that hinders the formation and growth of lymphatic endothelial cells (LECs). Recent findings of ectopic expression of LEC markers including Prospero Homeobox 1 (PROX1), a master regulator of lymphatic differentiation, and the vascular permeability marker Plasmalemma Vesicle Associated Protein (PLVAP), in certain glioblastoma and brain arteriovenous malformations (AVMs), has prompted investigation into their roles in cerebrovascular malformations, tumor environments, and blood-brain barrier (BBB) abnormalities. To explore the relationship between ectopic LEC properties and BBB disruption, we utilized endothelial cell-specific *Prox1* overexpression mutants. When induced during embryonic stages of BBB formation, endothelial *Prox1* expression induces hybrid blood-lymphatic phenotypes in the developing CNS vasculature. This effect is not observed when *Prox1* is overexpressed during postnatal BBB maturation. Ectopic *Prox1* expression leads to significant vascular malformations and enhanced vascular leakage, resulting in BBB disruption when induced during both embryonic and postnatal stages. Mechanistically, PROX1 downregulates critical BBB-associated genes, including *ß-catenin* and *Claudin-5*, which are essential for BBB development and maintenance. These findings suggest that PROX1 compromises BBB integrity by negatively regulating BBB-associated gene expression and Wnt/ß-catenin signaling.

## INTRODUCTION

The central nervous system (CNS), comprising both the brain and spinal cord, develops a specialized vascular network characterized by the presence of specialized endothelial cells (ECs) that constitute the blood-brain barrier (BBB) and the absence of lymphatic vasculature within the parenchyma. This barrier serves as a formidable separation blockade, dividing the CNS from the peripheral blood circulation (reviewed in(1–5)). The ECs comprising the BBB exhibit distinctive features compared to ECs in the other tissues: they possess continuous intercellular tight junction (TJ) proteins, lack fenestrations, and display minimal transcytosis activity(1–5). Furthermore, it is plausible that the absence of classical, highly permeable lymphatic capillaries, which are composed of lymphatic ECs (LECs) with discontinuous button-like junctions, impedes the induction of an immune response to CNS-derived antigens. This establishes the CNS parenchyma as an organ with immune-privileged status(6–8). Blood and lymphatic vasculature are closely associated in non-CNS tissues; however, the link between BBB integrity and lymphatic avascularity in the CNS parenchyma remains poorly understood.

LEC specification relies on the action of the homeobox transcription factor PROX1, which is necessary and sufficient to induce the LEC development program and repress the blood EC (BEC) development program in vitro and in vivo(9–15). Notably, LEC identity can be reprogrammed back into BEC identity by downregulating the expression of PROX1 during embryonic, postnatal, or adult stages(13). While the CNS parenchyma is considered an organ devoid of lymphatic vasculature, recent studies demonstrate that PROX1^+^ lymphatic vasculature develops an extensive network in the dura mater of meninges under the skull(16–19), and PROX1^+^ non-lumenized mural LECs, also called brain LECs or fluorescent granule perithelial cells, develop in the surface of zebrafish brain and mammalian leptomeninges(20–24). In several pathological conditions, including glioblastoma and brain arteriovenous malformations (AVMs), LEC markers including PROX1 are upregulated in ECs (25–27). Given that BBB integrity is often compromised in these glioblastoma and AVMs, these findings suggest a potential link between ectopic LEC marker expression and BBB disruption. Under normal physiological conditions, suppression of LEC properties may be essential for the development and maintenance of BBB in the CNS parenchyma. However, in pathological conditions, the ectopic upregulation of LEC markers might contribute to BBB disruption, thereby promoting disease progression.

In this study, we first analyzed publicly available single-cell RNA sequencing (scRNA-seq) data from human samples exhibiting impaired BBB integrity, including cases of AVMs, brain metastases, and glioblastoma tumors. Our analysis reveals upregulation lymphatic markers (*PROX1*, *LYVE1, FLT4*) in the CNS vasculature across these diseases associated with BBB dysfunction, alongside increased levels of *Plasmalemma Vesicle Associated Protein* (*PLVAP*), a factor commonly linked to endothelial permeability and BBB disruption. To explore the link between ectopic LEC differentiation in the CNS parenchyma and BBB disruption, we utilized a mouse model to express *Prox1* transgene, the master regulator of LEC development, in CNS ECs during BBB formation or maintenance. EC-specific overexpression of *Prox1* in mice results in significant alterations in the morphology and barrier function of the CNS vasculature.

Interestingly, endothelial *Prox1* expression induces a hybrid blood-lymphatic phenotype, characterized by the expression of both BEC markers and a subset of LEC markers, in the developing CNS vasculature when induced during primitive BBB formation at embryonic stages. However, such a hybrid blood-lymphatic phenotype is not observed when the *Prox1* expression is induced during the BBB maturation at postnatal stages.

However, endothelial *Prox1* expression promotes enhanced vascular leakage and BBB disruption when induced during both embryonic and postnatal stages. This vascular leakage is attributed to the downregulation of TJ proteins and the upregulation of transcytosis, underscoring the inhibitory effects of PROX1 on the BBB development and maintenance. Our in vitro experiments using brain ECs provide mechanistic insights into how PROX1 influences EC barrier functions: *Prox1* overexpression leads to a significant reduction in the expression of the TJ protein Claudin-5 and a destabilization of the actomyosin cytoskeleton, resulting in aberrant cell-cell junction formation. At the molecular level, PROX1 reduces the mRNA expression of BBB-associated genes, including *ß-catenin*, which is a critical signaling component for BBB development and maintenance. PROX1 disrupts BBB integrity through negative regulation of BBB-associated gene expression and Wnt/ß-catenin signaling. Collectively, our studies highlight the potential clinical impact of *Prox1* regulation in the CNS vasculature.

## RESULTS

### Lymphatic endothelial cell markers are upregulated in endothelial cells within brain tumors and vascular malformations

We analyzed publicly available single-cell RNA-seq (scRNA-seq) datasets from human glioblastoma (28–30), tumor metastases (31) and AVMs (32) to assess the expression of lymphatic endothelial cell (LEC) markers in endothelial cells (ECs) (Figure 1A and Supplemental Figure 1). After extracting ECs from three glioblastoma datasets and integrating them, we observed *PROX1* expression in ECs within the tumors, accompanying other lymphatic marker expressions (*LYVE1* and *FLT4*) (Figure 1B; Supplemental Figure 1, A-B). The tumor metastasis and AVM datasets each comprised disease (red) and control (blue) conditions, enabling comparisons between these states (Figure 1, C-D and Supplemental Figure 1, C-F). Examination of lymphatic marker genes revealed a pronounced increase in *PROX1* expression under disease conditions in both datasets. Additionally, we observed increased levels of *PLVAP*, which is commonly associated with endothelial permeability and BBB disruption(33–35), in ECs over all three disease conditions (Figure 1, B-D; Supplemental Figure 1, A-F). Also, downregulation of the BBB-associated markers (*CTNNB1* and *CLDN5*) was observed in disease conditions compared to control ECs (Supplemental Figure 1, D and F). These data suggest that abnormal differentiation from blood vessels to lymphatic vessels in the CNS may be connected to vascular permeability and BBB disruption observed in these conditions.

**Figure 1.**
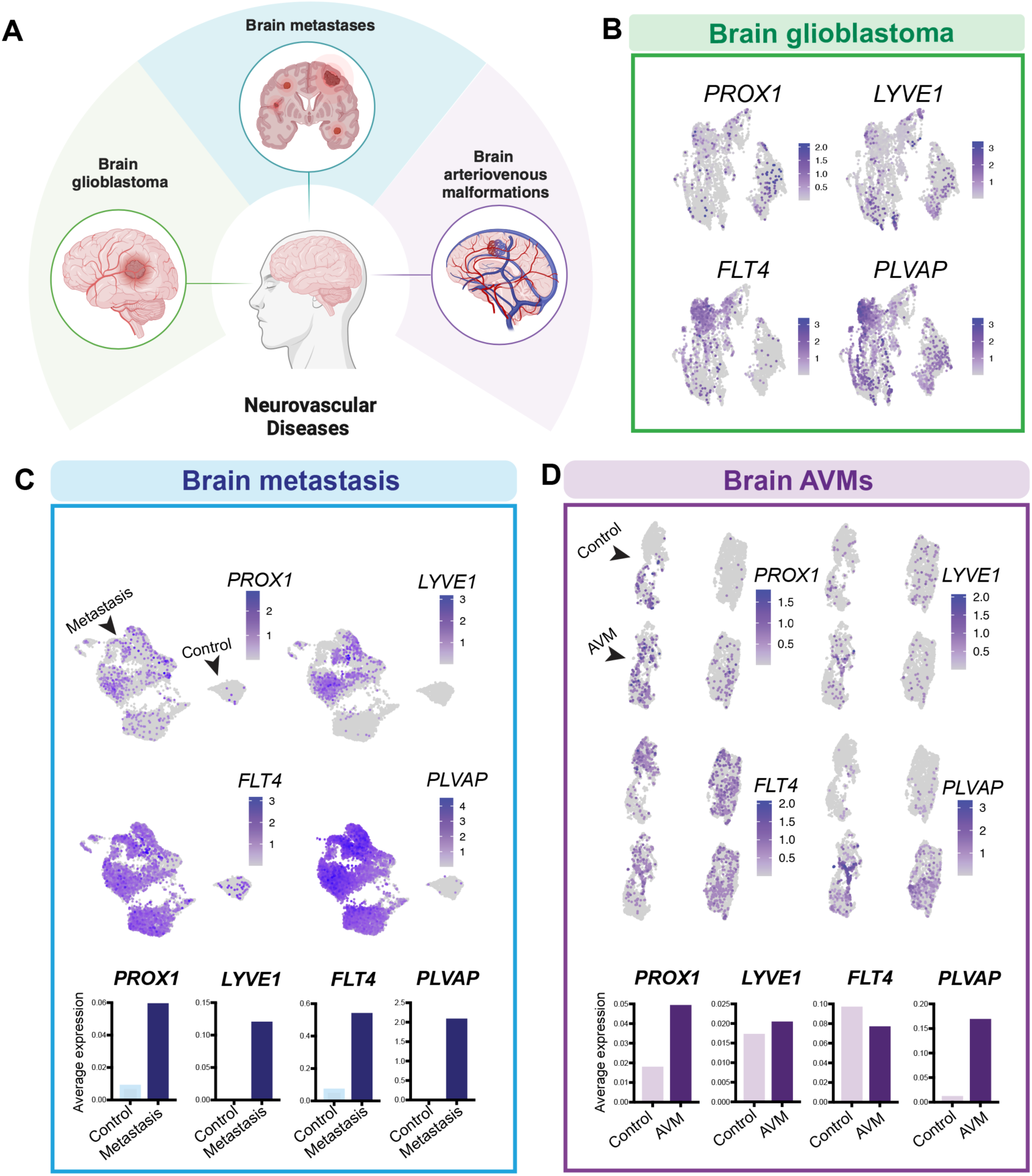
LEC markers are upregulated in ECs within human brain tumors and vascular malformations. **(A)** Schematic representation of human brain vascular diseases for publicly available single-cell RNA-seq (scRNA-seq) data analysis. scRNA-seq datasets for human glioblastoma (28–30), tumor metastasis (31) and brain arteriovenous malformations (AVMs) (32) were used. **(B)** UMAP plots of scRNA-seq data from glioblastoma datasets display endothelial cell (EC) clusters expressing LEC markers, including PROX1, LYVE1, and FLT4, along with the vascular permeability marker PLVAP. **(C)** UMAP plots and average gene expression charts of scRNA-seq data of control and metastatic brain tumor datasets display EC clusters expressing LEC markers, including PROX1, LYVE1, and FLT4, along with the vascular perme­ ability marker PLVAP **(D)** UMAP plots and average gene expression charts of scRNA-seq data of control and AVM datasets display EC clusters expressing LEC markers, including PROX1, LYVE1, and FLT4, along with the vascular permeability marker PLVAP

### Mouse CNS parenchyma lacks lymphatic vessels and does not exhibit temporal expression of PROX1 in its vasculature under physiological conditions

To investigate a potential link between ectopic LEC differentiation in the CNS parenchyma and BBB disruption, we turned to a mouse model to manipulate *Prox1* expression in the brain vasculature during primitive BBB formation at embryonic stages, or BBB maturation at postnatal stages. Given that the aforementioned scRNA-seq analysis from normal human brain samples revealed *PROX1* expression in a subset of brain ECs, we first examined PROX1 expression in the parenchymal vasculature of the mouse brain and spinal cord during embryonic or postnatal stages.

We performed high-resolution whole-mount imaging of mouse embryonic brains using the *Prox1-Gfp BAC* transgenic reporter(36), which allows visualization of PROX1-expressing cells with the green fluorescent protein (GFP). Since PROX1 is also expressed in neural progenitors and is recognized for its involvement in neuronal differentiation in the CNS(37), we defined PROX1-expressing ECs as those showing co-localization of GFP expression with both the pan-EC marker PECAM1 and the nuclear EC marker ERG. We also performed immunostaining using anti-PROX1 antibody to validate that the GFP signal corresponded to PROX1 expression. Section immunostaining of the *Prox1-Gfp* brain and spinal cord at embryonic stage (E)13.5 reveals that ERG^+^ EC-nuclei do not co-localize with PROX1 and Prox1-GFP, whereas there are numerous neural progenitors that are ERG-negative but positive for PROX1 and Prox1-GFP (Figure 2, A-C”; arrows indicate ERG^+^ EC-nuclei). Likewise, in the spinal cord parenchyma, ERG^+^ EC-nuclei do not co-localize with PROX1 and Prox1-GFP (Figure 2, D-E’, arrows). At E15.5, we did not observe any apparent co-localization of Prox1-GFP and the EC markers PECAM1 and ERG within a cluster of Prox1-GFP^+^ neural progenitors in the brain parenchyma (Figure 2F; arrows indicate ERG^+^/PECAM1^+^ ECs). The absence of PROX1 expression within the brain vasculature was confirmed during postnatal stages (Figure 2G). Of note, the combination of PECAM1 and LYVE1 allows us to corroborate the absence of classical lymphatic vessels (PECAM1^+^/LYVE1^+^/Prox1-GFP^+^) inside the brain parenchyma at postnatal stage (P)3, where only PECAM1^-^/LYVE1^+^/Prox1-GFP^-^ macrophages were found in the perivascular space (Figure 2, G-H, yellow arrowheads; Supplemental Figure 2, A-C). In contrast, PECAM1^+^/LYVE1^+^/Prox1-GFP^+^ lymphatic vessels were observed in both the meningeal layers and head skin vasculature (Figure 2, I-J, arrowheads; Supplemental Figure 2, D-H, arrows). Combined, this time-course analysis not only reaffirms the dearth of lymphatic vasculature within the CNS parenchyma but also underscores the absence of the lymphatic master regulator PROX1 in the CNS parenchyma ECs.

**Figure 2.**
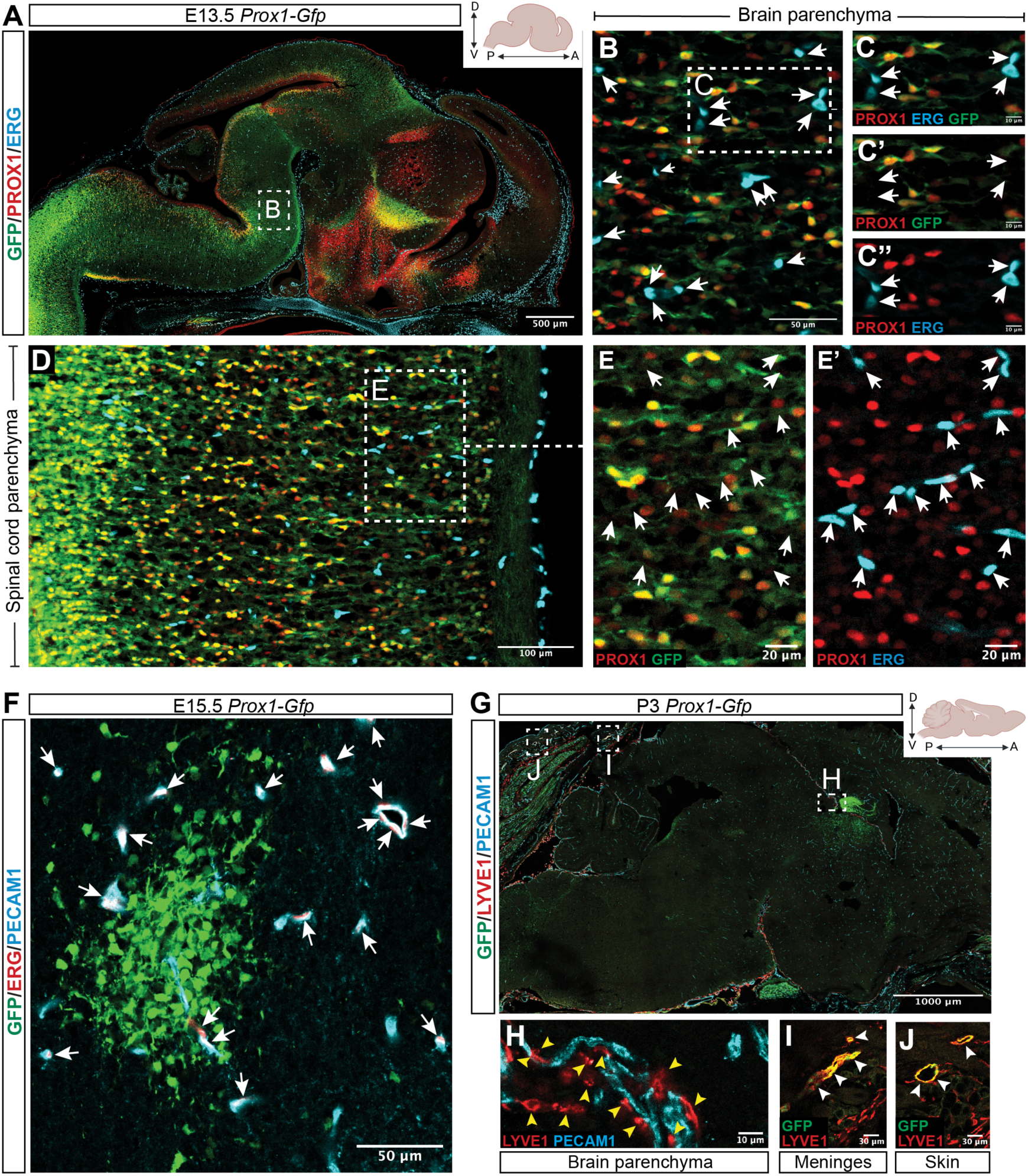
No temporal expression of Prox1 in the CNS vasculature. **(A-E)** A sagittal view of the brain (A-C) and spinal cord (D-E) parenchyma in E13.5 *Prox1-Gfp BAG* transgenic reporter embryos labeled with PROX1 (red) and ERG (cyan). The boxed regions in {A, B, and D) are magnified in {B, C-C”, and E-E’), respectively. Arrows indicate ERG+ EC nuclei in {B, C-C” and E-E’). These cells do not co-localize with PROX1 and *Prox1-GFP* Scale bars: 500 µmin (A), 100 µmin (D), 50 µmin (B), 20 µmin (E-E’), and 10 µmin (C-C”). **(F)** Brain parenchyma of E15.5 *Prox1-Gfp* embryos labeled with PECAM1 (cyan) and ERG (red). Arrows indicate ERG+/PECAM1+ ECs. These cells do not co-localize with *Prox1-GFP.* Scale bar: 50 µm. **(G-J)** A sagittal view of a postnatal brain section (P3) labeled with LYVE1 (red) and PECAM1 (cyan). The boxed regions in (G) are magnified in (H-J). Yellow arrowheads in (H) indicate LYVE1+/PECAM1-/Prox1-GFP­ macrophages. Arrowheads in (I and J) indicate PECAM1+/LYVE1+/Prox1-GFP+ lymphatic vessels in the meninges and skin, respectively. Scale bars: 1000 µmin (G), 30 µmin (I and J), and 10 µmin (H). The illustrations are created with BioRender.com.

### Endothelial *Prox1* expression leads to severe vascular abnormalities in the developing CNS vasculature

To address the relationship between PROX1 expression and BBB development/maintenance in a non-disease context, we generated a conditional *Prox1* overexpression mouse harboring the *loxP-STOP-loxP-Prox1* cassette in the *Rosa26* locus (*R26-LSL-Prox1* mice)(38), that allowed us to induce *Prox1* expression in a time- and cell type-specific manner (Supplemental Figure 3A). We further crossed them with the EC-specific *Cdh5-BAC-Cre^ERT2^*driver mice(39) to induce the *Prox1* transgene in ECs. Because previous studies have shown that primitive BBB becomes functional in the developing brain vasculature around E15.5(40), we opted to induce the *Prox1* transgene in *R26-LSL-Prox1* embryos (hereafter referred to as *Prox1^iEC-OE^*) through a tamoxifen administration at E13.5 and examine the resulting impact on brain vasculature development and BBB integrity at E16.5 (Figure 3A).

**Figure 3.**
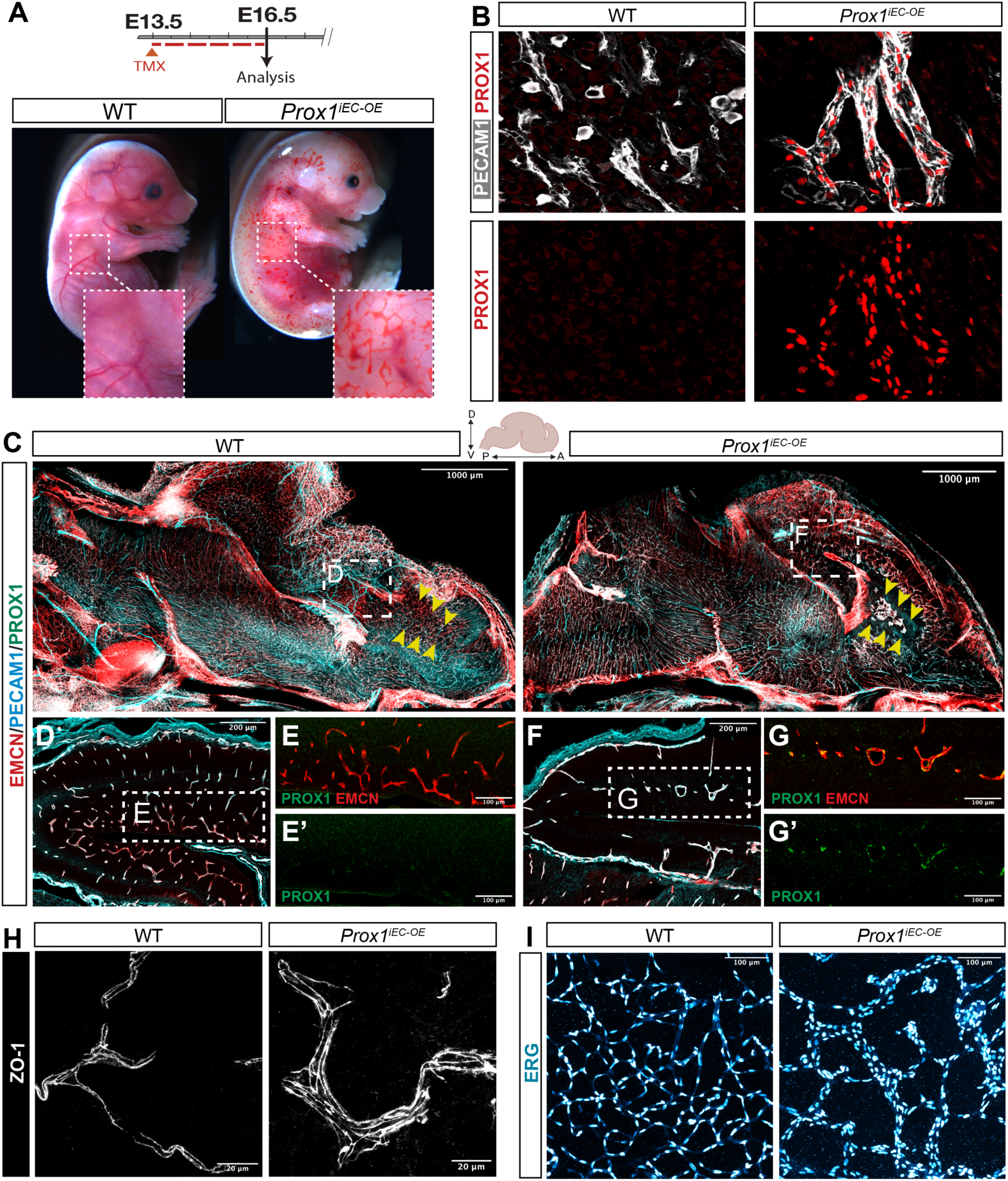
Endothelial *Prox1* expression induces vascular abnormalities in the developing CNS vasculature when induced during embryonic stages. (A) Diagram depicting EC-specific induction of *Prox1* expression at E13.5 and analysis of embryos at E16.5, and gross appearance of E16.5 *Prox1;cc.oc* mutant and their WT control littermate embryos. The boxed regions show blood-filled lymphatic vessels in the mutants when compared to control littermates. **(B)** Section immunostaining of E16.5 *Prox1K^0^E*mutant and their WT control littermate brains with antibodies to PROX1 (red) and PECAM1 (grey). Prox1 expression, detected with anti-PROX1 antibody (red), is significantly induced in PECAM1+ brain vasculature (grey) of E16.5 *Prox1i£C-oc* mutants compared to their WT control littermates. **(C-G)** A sagittal view of whole-mount immunostaining of E16.5 *Prox1’Ec-oE*mutant and their WT control littermate brains labeled with EMCN (red), PECAM1 (cyan) and PROX1 (green). The boxed regions in (C) are magnified in (D) as WT control and (F) as *Prox1’Ec-oc* mutant. The boxed regions in (D and F) are magnified in (E-E’ and G-G’), respectively. Yellow arrowheads in (C) indicate cortical vasculature. *Prox1,ec.oc* mutants exhibited PROX 1+IEMCN+ enlarged capillaries (F and G-G’) in comparison to their WT control littermates (D and E-E’). Scale bars: 1000 µmin (C), 200 µmin (D and F), and 100 µmin (E-E’ and G-G’). **(H-1)** Section immunostaining of E16.5 *Prox1^1^cc-oc* mutant and their WT control littermate brains with ZO-1 (H, grey) and ERG (I, hot cyan). *Prox11£c-oc* mutants exhibited enlarged **capillaries with an increased number of ECs in comparison to their WT control littermates. Scale bars: 100 µm in (I) and 20 µm in (H). The illustrations are created** with BioRender.com.

The resulting *Prox1^iEC-OE^* mutant embryos exhibited pronounced edemas, hemorrhagic manifestation, and blood-filled lymphatics in the skin (Figure 3A; Supplemental Figure 3B). Moreover, the mutants exhibited embryonic lethality within 72 hours following the *Prox1* transgene induction in ECs (Supplemental Figure 3B). We validated the efficient induction of the *Prox1* transgene in PECAM1^+^ ECs of the brain vasculature in *Prox1^iEC-OE^* mutant embryos, whereas *Prox1* expression was absent in PECAM1^+^ ECs in their *wild-type* (WT) control littermates (Figure 3B). A sagittal overview reveals significant disparities in the brain vasculature between *Prox1^iEC-OE^* mutant embryos and their WT control littermates, notably in the cerebral cortex region where abnormal enlarged vessels are present, while capillary density is reduced in the mutants (Figure 3, C-G, yellow arrowheads; Supplemental Figure 3, C-H). Immunostaining with antibodies to the adherent junction marker ZO-1 and the nuclear EC marker ERG reveals the formation of thick capillaries due to an augmented number of ECs in *Prox1^iEC-OE^* mutant embryos, as compared to their WT control littermates (Figure 3, H-I).

We next investigated whether *Prox1* expression induces a LEC fate in the vasculature of *Prox1^iEC-OE^* mutant embryos. We first examined the expression of the classical LEC marker LYVE1 in the vasculature of *Prox1^iEC-OE^* mutant embryos and their WT control littermates. We observed a substantial increase in PECAM1^+^/LYVE1^+^ lymphatic vessels in the trunk vasculature of the mutant embryos compared to their WT control littermates (Figure 4A, arrows). In contrast, we did not detect any LYVE1-expressing ECs in the brain vasculature of either the mutant embryos or their WT control littermates (Figure 4B). Quantitative validation of these findings was achieved through flow cytometry analysis (Supplemental Figure 4, A-B): PECAM1^+^/LYVE1^+^ LECs were not detectable in both the mutant and control brain (comprising 0% of brain ECs), whereas the mutant skin exhibited a significant increase in the proportion of LYVE1^+^/PECAM1^+^ LECs (from 5% to 30% of skin ECs) alongside a concurrent decrease in LYVE1^-^/PECAM1^+^ BECs (from 95% to 70% of skin ECs). These data suggest that consistent with the established propensity of PROX1 function to evoke lymphatic differentiation in the developing vasculature, endothelial *Prox1* expression induces the differentiation of BECs into LECs in the skin vasculature. In contrast, in the brain vasculature, *Prox1* does not induce conventional LECs. While *Prox1* induces significant remodeling in the brain parenchymal vasculature, characterized by the rapid development of enlarged vessels and thicker capillaries, particularly in the cerebral cortex region, it appears that *Prox1* expression alone is insufficient to induce conventional LECs expressing the classical LEC markers such as LYVE1 (Figure 4B) and podoplanin (PDPL, data not shown).

**Figure 4.**
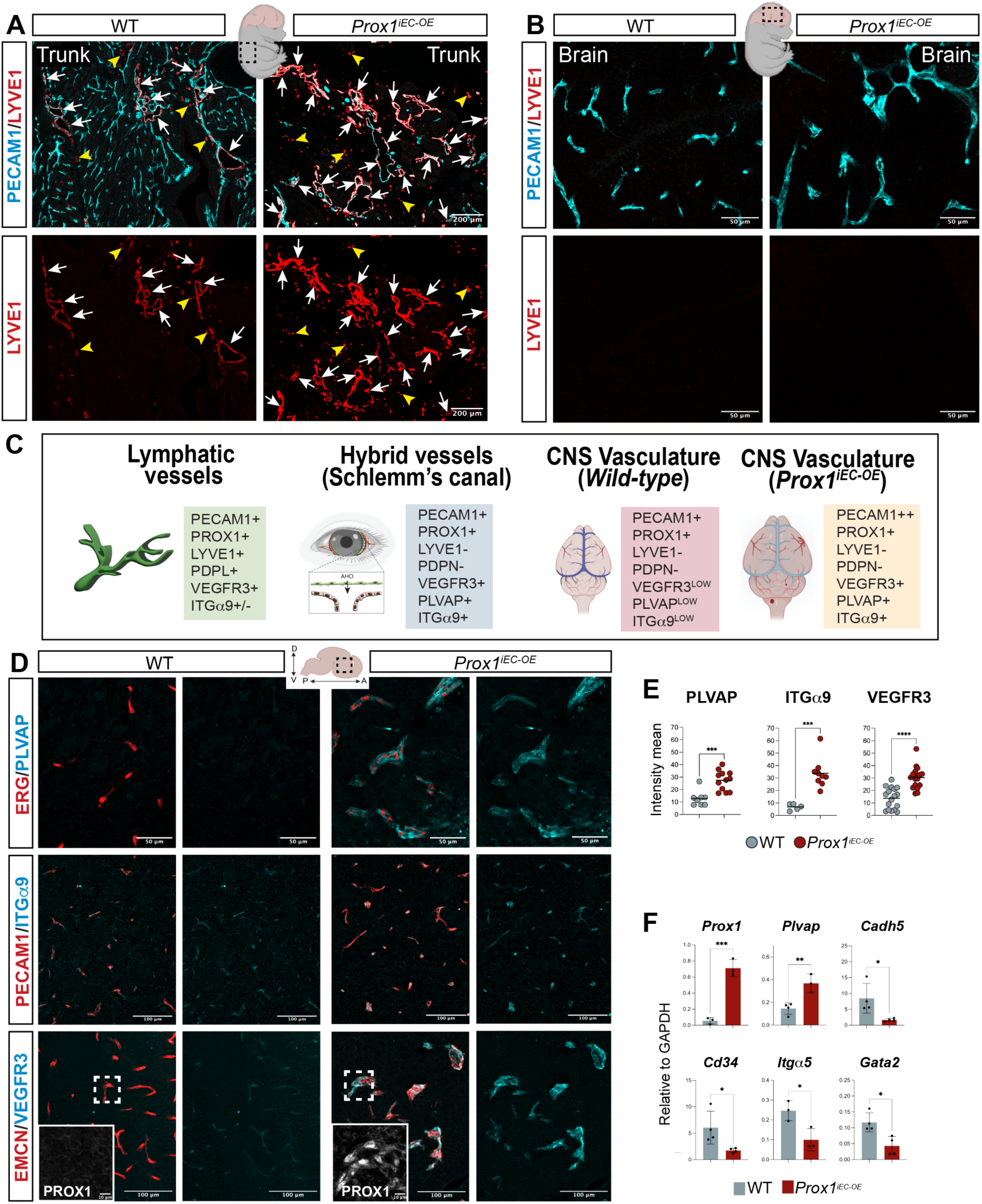
Endothelial *Prox1* expression induces a hybrid blood-lymphatic phenotype in the developing CNS vasculature. **(A-8)** Section immunostaining of trunk (A) and brain (B) from E16.5 *Proxt’ec-OE*mutant and their WT control littermate embryos with PECAM1 (cyan) and LYVE1 (red). **Arrows indicate PECAM1+/LVYE1+ lymphatic vessels, while yellow arrowheads indicate PECAM1-/LVYE1+ macrophages.** *ProxtfEC-oE* **mutants exhibited enhanced** lymphatic differentiation to form PECAM1+/LVYE1+ lymphatic vasculature in the trunk in comparison to their WT control littermates. In contrast, both *Prox1•ee-oe* mutant and their WT control littermates did not exhibit conventional PECAM1+/LVYE1+ lymphatic vasculature in the brain. Scale bars: 200 µmin (A) and 50 µmin (B). **(C)** Schematic illustrations indicating EC markers expressed in conventional lymphatic vessels, hybrid vessels in the Schlemm’s canal, and CNS vessels in WT and *Proxt’ec-oe* mutants. **(D-E)** Section immunostaining of E16.5 *Prox1.ec-0e* mutant and their WT control littermate brains with PLVAP (cyan), ITGa9 (cyan) and VEGFR3 (cyan), together with ERG (red), PECAM1 (red), EMCN (red), respectively. The sections labeled with VEGFR3 (cyan) and EMCN (red) are additionally stained with PROX1 (grey in the magnified images). (E) Quantifications of the fluorescence intensity mean for PLVAP, ITGu9 and VEGFR3 in the brain vasculature using lmaris software. Each dot corresponds to random fields of view from at least 3 different WT control and mutant embryos. Data are shown as mean±SEM. Scale bars: 100 µm and 50 µmin (D). (F) Relative mRNAexpression levels of *Prox1* and *Plvap* together with BEC markers such as *Cadh5, Cd34, ltga5,* and *Gata2* in FACS-isolated brain ECs from E16.5 *Proxt’ec-oe*mutant and their WT control littermate brain. Bar graphs show mean normalized expression ±SEM; n=3-4 biological samples obtained from **FAGS-isolated brain ECs from individual experiments.,.,.. p<0.001, “’”’*p<0.0005, “’”’”’*p<0.0001, as determined by unpaired t-test. The illustrations are created with** BioRender.com.

Of note, given our use of the EC-specific *Cdh5-BAC-Cre^ERT2^*driver mice to induce the *Prox1* transgene in ECs, we observed abnormalities in the lymphatic vasculature in peripheral tissues. For instance, whole-mount immunostaining of limb skin and heart ventricles revealed aberrant branching of lymphatic vessels in *Prox1^iEC-OE^* mutant embryos (Supplemental Figure 4, C-E). As previously described(41), LYVE1^+^/PECAM1^+^ cardiac lymphatic vessels extend inferior on both the ventral and dorsal surfaces of the heart ventricle in the WT control littermates (Supplemental Figure 4D, arrows). Notably, some of these lymphatic vessels branch closely to EMCN^+^/PECAM1^+^ large-diameter coronary veins on the dorsal surface of the heart ventricle. In contrast, the ventral surface of the mutant heart ventricle exhibited blood-filled lymphatic vasculature, while the dorsal surface showed abnormal lymphatic structures (Supplemental Figure 4E).

Additionally, the mutants exhibited underdeveloped coronary vasculature, characterized by the absence of large-diameter coronary arteries (Supplemental Figure 4, D-E, PECAM1^+^, arrowheads) and veins (Supplemental Figure 4, D-E, EMCN^+^, yellow arrowheads). These findings suggest that endothelial *Prox1* expression leads to abnormal coronary and cardiac lymphatic vasculature in the developing heart ventricles.

### Endothelial *Prox1* expression induces a hybrid blood-lymphatic phenotype in the developing CNS vasculature

In light of the recent discovery of Schlemm’s canal in the eye, which is a specialized ring-shaped vasculature at the periphery of the cornea and has ECs having BEC and LEC characteristics, including the expression of BEC markers and a subset of LEC makers(42–44), we proceeded to examine whether *Prox1* expression induces a hybrid blood-lymphatic phenotype in the brain vasculature. Schlemm’s canal ECs manifest the expression of BEC markers including PECAM1, endomucin (EMCN), CD34, VE-Cadherin (Cdh5), and Tie2, and the expression of LEC markers such as PROX1, VEGFR3, and ITGα9. The classical LEC markers LYVE1 and PDPL are absent in Schlemm’s canal ECs (Figure 4C). Additionally, plasmalemma vesicle-associated protein (PLVAP), a component of endothelial fenestrae that regulates basal permeability(33–35), is highly expressed Schlemm’s canal ECs (Figure 4C).

In the brain vasculature of WT control littermates, the expression of PLVAP, VEGFR3, and ITGα9 is scarcely detectable in ECs (Figure 4, D-E; overlap with the pan-EC markers ERG and PECAM1, or the pan-capillary EC marker EMCN). In contrast, in the brain vasculature of *Prox1^iEC-OE^* mutant embryos, the expression of these markers is substantially upregulated compared to their WT control littermates (Figure 4D; quantification in Figure 4E). At the mRNA level, brain ECs isolated through fluorescence-activated cell sorting (FACS) from *Prox1^iEC-OE^* mutant embryos demonstrate increased expression of *Plvap* compared to their WT control littermates (Figure 4F). Although the expression of BEC markers such as *VE-Cadherin*/*Cdh5*, *Cd34*, *Itg*α*5,* and *Gata2* is partially reduced in FACS-isolated brain ECs from *Prox1^iEC-OE^* mutant embryos compared to their WT control littermates (Figure 4F), it is evident that endothelial *Prox1* expression does not completely reprogram BECs to LECs in the brain vasculature. Taken together, this evidence shows that *Prox1* induces a hybrid blood-lymphatic phenotype in the brain vasculature, reminiscent of Schlemm’s canal ECs in the eyes, with the expression of BEC (PECAM1^+^/PLVAP^+^) and LEC (PROX1^+^/VEGFR3^+^/ITGα9^+^) markers.

### Endothelial expression of *Prox1* disrupts primitive blood-brain barrier formation in the developing CNS vasculature

CNS ECs express the tight junction (TJ) protein Claudin-5 (CLDN5) as a marker for BBB, while PLVAP, inductive of high-permeability vasculature, is normally absent from these cells(5, 45). In regions of the brain where the BBB is compromised, there is a reduction in CLDN5 expression and an induction of PLVAP(5, 46). We then investigated whether the acquisition of such a hybrid blood-lymphatic phenotype in the CNS vasculature of *Prox1^iEC-OE^* mutant embryos might affect the development and integrity of the BBB.

Immunostaining with antibodies to the TJ marker CLDN5 and the pan-EC marker ERG clearly demonstrates a reduction in CLDN5 expression in the brain vasculature of *Prox1^iEC-OE^* mutant embryos as compared to their WT control littermates (Figure 5A; quantification in Figure 5B). This reduction indicates impaired TJ assembly among cerebral ECs, suggesting a defect in barrier integrity. Moreover, Ter119^+^ blood cell extravasation was observed in *Prox1^iEC-OE^* mutant brains (Supplemental Figure 5, A-C, arrows). These data indicate a potential compromise in the BBB. To further address whether the mutant brains exhibited compromised barrier function, we performed a tracer leakage assay in E16.5 *Prox1^iEC-OE^* mutant embryos and their WT control littermates when the primitive BBB becomes functional(40). We harvested embryos and performed an intracardial injection of a 3kDa fluorescent tracer, Dextran Texas-Red (Figure 5C). Whole brain images and subsequent immunostaining of sagittal brain samples reveal extensive BBB leakage in *Prox1^iEC-OE^* mutant embryos (Figure 5, D-H; quantification in Figure 5I; Supplemental Figure 5, D-F): The injected dextran tracer remained entirely within PECAM1^+^ vasculature of WT control littermates (Figure 5, D-F and Supplemental Video 1), while severe BBB leakage was observed in the mutant embryos, particularly within the cerebral cortex (Figure 5, D-H and Supplemental Video 2). These findings suggest that the endothelial *Prox1* expression disrupts primitive BBB formation in the developing CNS vasculature.

**Figure 5.**
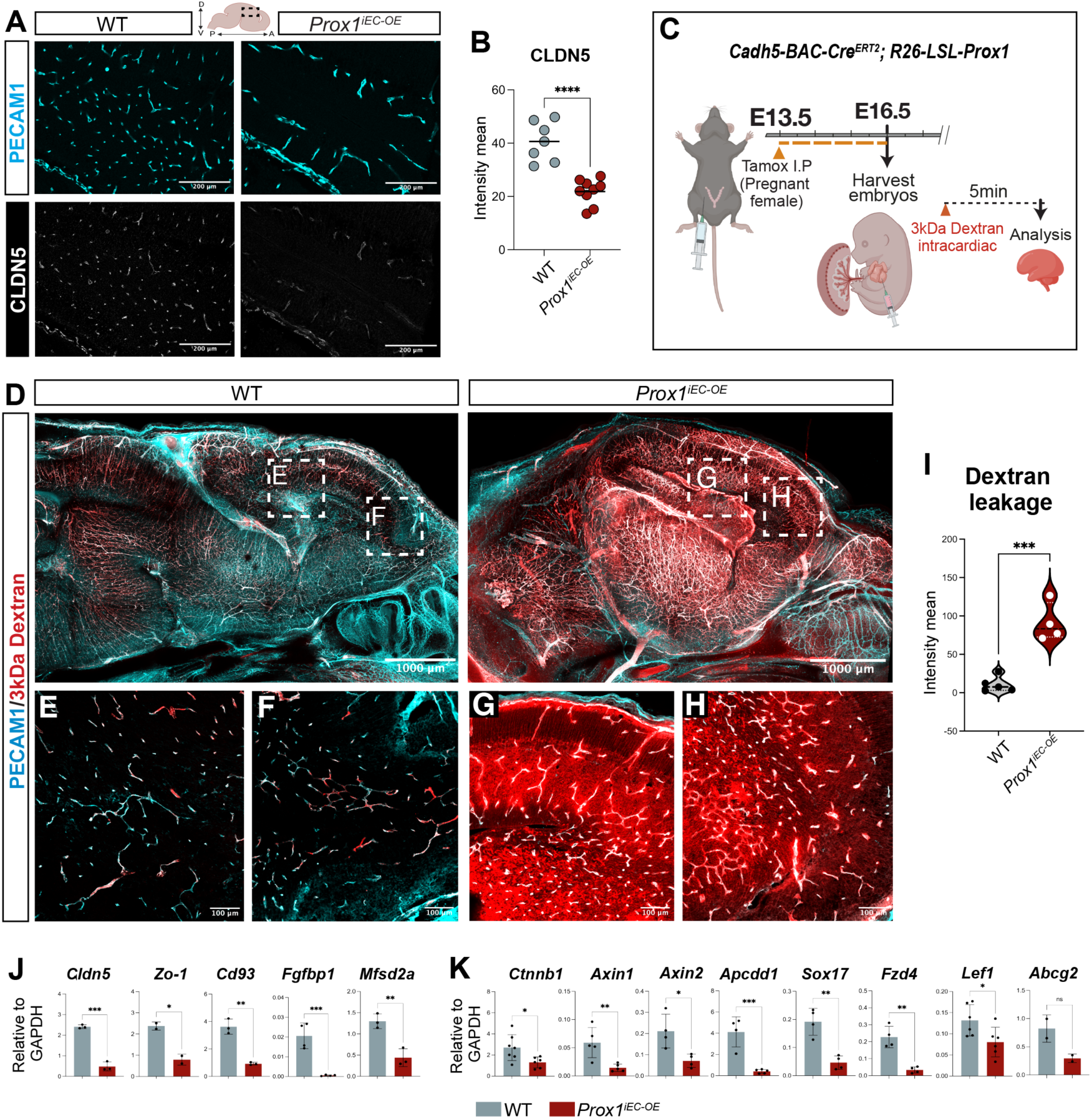
Endothelial *Prox1* expression disrupts the primitive blood-brain barrier formation in the developing CNS vasculature. **(A-B)** Section immunostaining of E16.5 *Prox1^1^ec-oe* mutant and their WT control littermate brains with PECAM1 (cyan) and CLDN5 (grey). The expression of CLDN5 is downregulated in the brain vasculature of Prox1’ec-0e mutants in comparison to their WT control littermates. (B) Quantifica­ tions of the fluorescence intensity mean for CLDN5 in the brain vasculature using lmaris software. Each dot corresponds to random fields of view from at least 3 different WT control and mutant embryos. Data are shown as mean±SEM. ****p<0.0001, as determined by unpaired I-test. Scale bars: 200 µmin (A). (C) Diagram depicting EC-specific induction of *Prox1* expression at E13.5 and vascular permeability analysis of embryos at E16.5. (D-l) A sagittal view of whole-mount imaging of E16.5 *Prox1’ec-oe*mutant and their WT control littermate brains with 3kDa Dextran Texas-Red tracer (red) and PECAM1 (cyan). Boxed regions in (D) are magnified in (E and F) as WT control littenmates and (G and H) as *Prox1’ecoe* mutants. The mutant brains exhibited extensive BBB leakage. (I) Quantification of Dextran Texas-Red tracer extravasation outside of the brain vasculature in WT control (n=5 individual brains, showing the average of 4 different fields of view) and mutant brains (n=4 individual brains, showing the average of4 different fields of view). ***p<0.0005, as determined by unpaired I-test Scale bars: 1000 µmin (D) and 100 µmin (E-H). **(J-K)** Relative mRNA expression levels of BBB-related genes such as *Cldn5, Zo-1, Cd93, Fgfbp1,* and *Mfsd2a* in (J), and 11-catenin and its target genes such *Ctnnb1, Axin1, Axin2, Apcdd1, Sox17, Fzd4, Left,* and *Abcg2* in (K), in FAGS-isolated brain ECs from E16.5 *Prox1^1^ec-oe* mutant and their WT control littermate brain. Graphs show mean normalized expression ±SEM; n=3-5 experiments obtained from 4 individual FAGS experiments. *p<0.05, “” p<0.001 ‘*’p<0.0005, ‘*”p<0.0001, as determined by unpaired I-test. The illustrations are created with BioRender.com.

We next assessed the mRNA expression of BBB markers in FACS-isolated ECs from *Prox1^iEC-OE^* mutant brains and their WT control littermates. We observed a decrease in the expression of TJ markers *Cldn5* and *Tjp1/ZO-1* in *Prox1^iEC-OE^* mutant embryos compared to their WT control littermates (Figure 5J). We also observed a decrease in the expression of recently identified BBB-related genes, such as *Cd93*(*47*) and *Fgfpd1*(*48*) in the mutant embryos compared to their WT control littermates (Figure 5J). Additionally, we found a reduction in the expression of the lipid transporter *Mfsd2a*, which plays an essential role in limiting caveolin-dependent transcytosis in BBB ECs(40, 49–51) in the mutant embryos compared to their WT control littermates. Furthermore, the expression of *Pten*, which serves as an upstream regulator of the Mfsd2a-transcytosis axis(51), was also downregulated in the mutant embryos (Supplemental Figure 5G). This finding suggests a potential upregulation of transcytosis in addition to an impaired TJ in the mutant embryos (Figure 5J). Given that Wnt/ß-catenin signaling is known to regulate many BBB genes including *Cldn5*, *Plvap*, and *Mfsd2a*(1, 4, 5), we observed a decrease in the expression of *Ctnnb1/ß-catenin* as well as several effector and target genes associated with Wnt/ß-catenin signaling in the mutant embryos compared to their WT control littermates (Figure 5K and Supplemental Figure 5G). These results indicate that the endothelial *Prox1* expression leads to a significant downregulation of Wnt/ß-catenin signaling in the developing CNS vasculature.

Pericyte-EC association is essential for the formation of a functionally effective BBB(52, 53). Thus, barrier defects in *Prox1^iEC-OE^* mutant embryos could be due to altered pericyte coverage of capillaries. However, immunostaining with antibodies to the pericyte markers NG2 and PDGFRß, in combination with PECAM1, reveals pericyte coverage of enlarged capillaries in the brain vasculature of *Prox1^iEC-OE^* mutant embryos (Supplemental Figure 5, H-K, arrows). Indeed, FACS analysis reveals a comparable number of CD140b(PDGFRß)^+^/CD31(PECAM1)^-^ pericytes in both *Prox1^iEC-OE^* mutant embryos and their WT control littermates, exhibiting a similar maximal fluorescence intensity (MFI) corresponding to the expression of the pericyte marker CD140b (Supplemental Figure 5, L-M). Of note, immunostaining with the anti-NG2 antibody also labels oligodendrocytes (NG2^+^/PDGFRß^-^) (Supplemental Figure 5, H-K, yellow arrowheads), and we observed an increased association between oligodendrocytes and capillaries (Supplemental Figure 5, I and K, yellow arrowheads) in *Prox1^iEC-OE^* mutant embryos compared to their WT control littermates. Given that previous studies have reported the expression of Wnt7a/b ligands for canonical Wnt/ß-catenin signaling(54–56) by oligodendrocytes, in addition to astroglia and neurons, these findings suggest that oligodendrocytes may play a role in repairing BBB disruption.

### Postnatal induction of *Prox1* leads to blood-brain barrier breakdown

The observation that the endothelial *Prox1* expression during primitive BBB formation leads to the BBB disruption prompted us to investigate whether PROX1 itself exerts any influence on BBB function after it has already formed and matured, even in the absence of LEC differentiation in the CNS parenchyma. To address this question, we opted to induce the *Prox1* transgene through tamoxifen administration during BBB maturation at postnatal stage (P)7 and examine the resulting impact on BBB integrity. We performed a tracer leakage assay in P10 *Prox1^iEC-OE^* mutant mice and their WT control littermates: We performed an intraperitoneal injection (I.P.) of a 3kDa Dextran Texas-Red or a 1kDa Alexa Fluor 555-Cadaverine (Figure 6A). Brightfield whole brain images show enlarged vessels in *Prox1^iEC-OE^* mutant brains (Figure 6B). Whole-mount immunostaining and tissue clearing of sagittal brain samples with antibodies to the EC markers PECAM1 or EMCN reveals extensive BBB leakage in *Prox1^iEC-OE^* mutant brains (Figure 6, D-F; quantification in Figure 6C). Severe BBB leakage was observed within the cerebellum of the mutant mice (Figure 6, E-E’ and F-F’). Subsequent section immunostaining of the cerebellum clearly demonstrates that the dextran tracer leaks out of vessels in the mutant mice (Figure 6G). We also observed similar leakage in the 1kDa Alexa Fluor 555-cadaverine tracer (Supplemental Figure 6B; quantification in Supplemental Figure 6D). These results show that the endothelial *Prox1* expression disrupts barrier function in the postnatal CNS vasculature.

**Figure 6.**
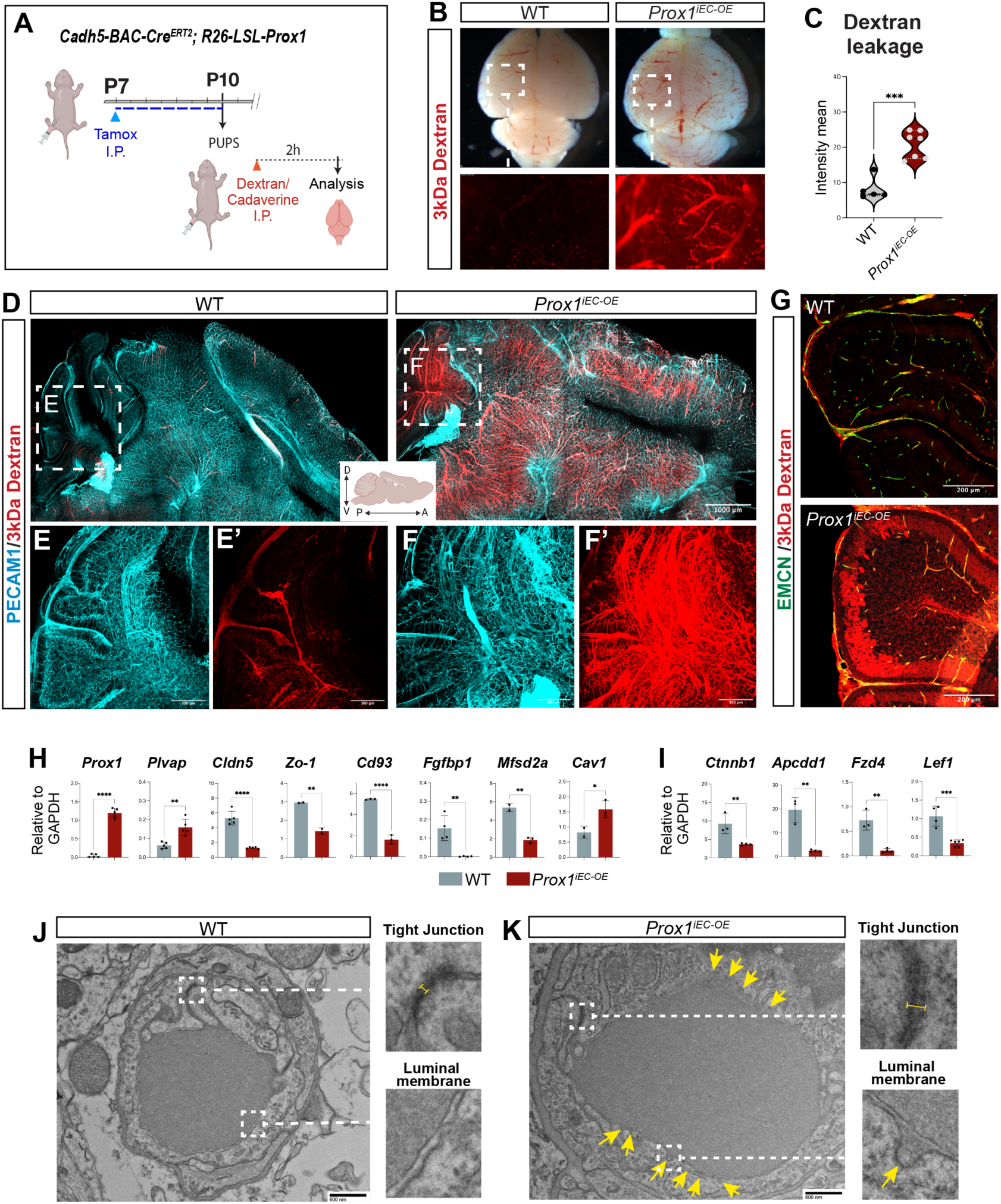
Postnatal induction of *Prox1* disrupts the blood-brain barrier. **(A)** Diagram depicting EC-specific induction of *Prox1* expression at P7 and vascular permeability analysis of pups at P10. **(B-C)** Gross appearance of P10 *Prox/iEC-oe* mutant and their WT control littermate brains with 3kDa Dextran Texas-Red tracer (red). Boxed regions are magnified in the lower panels. (CJ Quantification of Dextran Texas-Red tracer outside of the brain vasculature in WT control (n=5 individual brains, showing the average of 4 different fields of view) and mutant brains (n=7 individ­ ual brains. showing the average of 4 different fields of view). …p<0.0005, as determined by unpaired I-test. (D-F) A sagittal view of whole-mount imaging of P10 *Prox/iEC-OE* mutant and their WT control littermate brains with 3kDa Dextran Texas-Red tracer (red) and PECAM 1 (cyan). Boxed regions in (D) are magnified in (E-E’) as WT control littermates and (F-F’) as *ProxfEc-oe* mutants. The mutant brains exhibited extensive BBB leakage in the cerebellum. Scale bars: 1000 µm in (D) and 300 µmin (E-F). (G) Section immunostaining of P10 *Prox/;Ec-oe* mutant and their WT control littermate cerebrum with 3kDa Dextran Texas-Red tracer (red) and EMCN (green). Scale bars: 200 µm. (H-I) Relative mRNA expression levels of *Proxt* and *Plvap* together with BBB-related genes such as *Cldn5, Zo-1, Cd93, Fgfbp1, Mfsd2a,* and *Cavt* in (H) and P..-catenin and its target genes such *Ctnnb1, Apcdd1, Fzd4,* and *Left* in (I), in FAGS-isolated brain ECs from P10 *Proxt•Ec-oE*mutant and their WT control littermate brain. Graphs show mean normalized expression ±SEM; n=3-4 biological samples obtained from FACS-isolated brain ECs from individual experi­ ments… p<0.001 … p<0.0005, p<0.0001, as determined by unpaired t-test. **(J-K)** Representative transmission electron microscopy (TEM) images of brain capillaries in P10 *Prox1”e-oE* mutant and their WT control littermate brain. The boxed regions show tight junctions and luminal membranes are magnified in the right panels. Yellow arrows indicate endothelial vesicles in (K). Scale bars: 600nm in (J and K). The illustrations are created with BioRender.com.

We next investigated whether the *Prox1* expression impacts capillary network and BBB integrity. While the brain of the WT control littermate featured a dense capillary network, the mutant brain exhibited abnormally enlarged vasculature with reduced vascular density and larger caliber vessels (Supplemental Figure 6A). However, we did not detect any significant change in the mRNA expression of BEC markers such as *VE-Cadherin*/*Cdh5*, *Cd34*, *Itg*α*5,* and *Gata2* between *Prox1^iEC-OE^*mutants and their WT control littermates (Supplemental Figure 6F). Moreover, we also did not observe a hybrid blood-lymphatic phenotype in the postnatal brain vasculature of *Prox1^iEC-OE^* mutants: we did not find upregulation of LEC markers such as VEGFR3 and ITGα9, as was observed in the developing brain vasculature (Supplemental Figure 6, G-H). These findings suggest that the *Prox1* does not induce a hybrid blood-lymphatic phenotype in the postnatal brain vasculature.

Since impaired barrier function correlates with impaired TJ proteins, we observed a reduction in the expression of CLDN5 in the brain vasculature of *Prox1^iEC-OE^* mutants as compared to their WT control littermates (Supplemental Figure 6B; quantification in Supplemental Figure 6C). Supporting this observation, we also found a decrease in the mRNA expression of BBB markers, such as *Cldn5*, *Tjp1/ZO-1*, *Cd93*, *Fgfbp1,* and *Mfsd2a,* and an increase in the expression of *Plvap* and *Caveolin-1/Cav1,* in *Prox1^iEC-OE^* mutants compared to their WT control littermates (Figure 6H). These findings demonstrate that the endothelial *Prox1* expression disrupts barrier integrity in the postnatal CNS vasculature.

Given that EC ß-catenin signaling is known to maintain the BBB state(46, 57–60), we observed a decrease in the mRNA expression of *Ctnnb1/ß-catenin,* as well as several effector and target genes associated with Wnt/ß-catenin signaling in the mutants compared to their WT control littermates (Figure 6I and Supplemental Figure 6E). Taken together with the findings from the analysis of the developing CNS vasculature, these data show that the endothelial *Prox1* expression significantly downregulates Wnt/ß-catenin signaling in both developing and postnatal CNS vasculature.

Recent observations indicating that Wnt/ß-catenin signaling activates *Mfsd2a* to limit caveolae-mediated transcytosis in CNS ECs(50, 51, 57, 61) prompted us to investigate whether the *Prox1* expression affects both transcellular and the aforesaid paracellular permeability in the postnatal CNS vasculature. Through transmission electron microscopy (TEM) analysis, we first observed an enlarged capillary lumen in *Prox1^iEC-OE^* mutants compared to their WT control littermates (Figure 6, J and K). Secondly, we verified an increased gap in TJ and an increased number of transcellular vesicles in ECs of the mutants in comparison to their WT control littermates (Figure 6K, yellow arrows). These data indicate that the endothelial *Prox1* expression induces BBB breakdown by enhancing both paracellular and transcellular permeability in the postnatal CNS vasculature.

### Endothelial *Prox1* expression induces abnormal tight junctions by repressing *Claudin-5* expression and destabilizing actin filaments in brain endothelial cells

We next explored how PROX1 disrupts EC barrier functions. To address this question, we turned to in vitro culture experiments using a mouse brain EC line, bEnd.3 cells, known for its brain EC-specific characteristics, including the maintenance of neural stem cells(62). Importantly, previous studies demonstrated that Wnt/ß-catenin signaling upregulates the expression of *Mfsd2a*, while downregulating the expression of *Cav1* and *Plvap* in cultured bEnd.3 cells(57). Given that endogenous PROX1 expression was not detectable in bEnd.3 cells, we introduced the *Prox1* or *Gfp* transgene into the cells using a lentiviral system and subsequently cultured these infected cells until they formed confluent monolayers (Supplemental Figure 7, A-B). Most of the bEnd.3 cells expressing *Prox1* exhibited discontinuous cell-cell junctions and an enlarged cell shape, as determined with ZO-1 immunostaining, whereas the bEnd.3 cells expressing *Gfp* showed continuous yet reticular cell-cell junctions without altered cell shape (Figure 7, A-B; three representative images for each bEnd.3 cells expressing *Prox1* or *Gfp*). Since EC junctions are tightly regulated by actin cytoskeleton(63), we observed a significant reduction in the intensity of F-Actin (Figure 7, A-B and C-D) and phospho-myosin light chain 2 (p-MLC2), a downstream target of the RhoA/ROCK pathway that regulates actin stress fiber contraction and cytoskeleton remodeling, in the bEnd.3 cells expressing *Prox1* (Figure 7, C-D). These observations suggest that the *Prox1* expression leads to abnormal organization and a relaxation of actin stress fibers, resulting in the formation of enlarged cell shape and abnormal cell-cell junctions. Indeed, the bEnd.3 cells expressing *Prox1* not only exhibited discontinuous cell-cell junctions but also a marked reduction in CLDN5 expression. In contrast, the bEnd.3 cells expressing *Gfp* showed colocalization of ZO-1 and CLDN5 in continuous cell-cell junctions (Figure 7, E-F; quantification in Figure 7G). Of note, we also observed abnormal cell-cell junctions in most primary rat brain microvascular ECs expressing *Prox1* (RBMVECs) (Supplemental Figure 7, C-D). Collectively, these in vitro studies present compelling evidence of abnormal TJs because of the endothelial *Prox1* expression in brain ECs. Interestingly, *Prox1* also influences the actin cytoskeleton, promoting disorganized and relaxed actin fibers, which in turn result in less polarized and enlarged ECs. These observations may correspond to the formation of enlarged vessels and thicker capillaries in the CNS vasculature of *Prox1^iEC-OE^* mutant mice.

**Figure 7.**
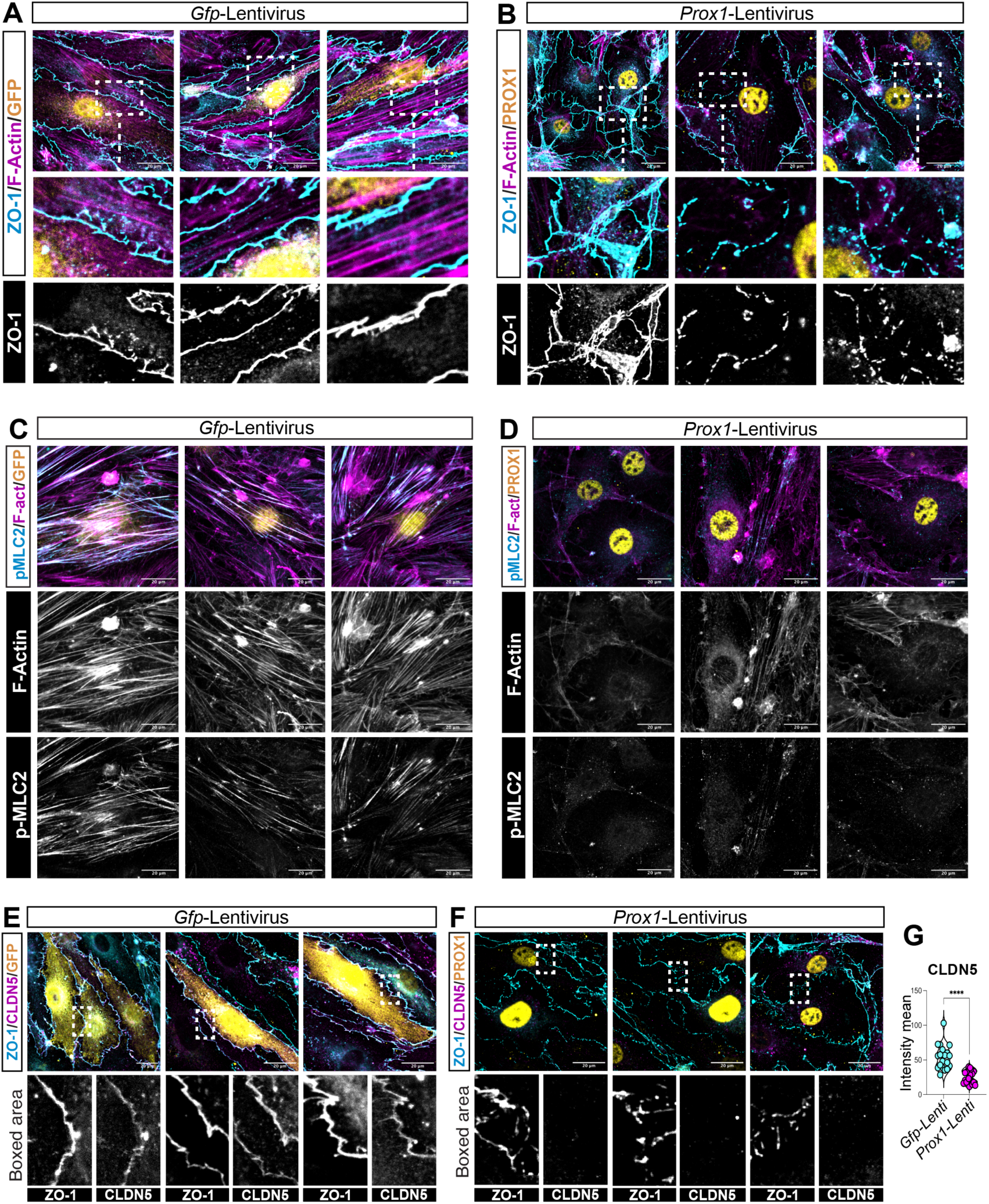
*Prox1* expression induces abnormal tight junctions by repressing *C/audin-5* expression and destabilizing actin filaments in cultured bEnd.3 cells. (A-B) Three representative images of cultured bEnd.3 mouse brain ECs expressing *Gfp* or *Prox1* labeled with ZO-1 (cyan or grey) and F-Actin (magenta) together with GFP (yellow) or PROX1 (yellow). respectively. Boxed regions are magnified **in** the lower panels. bEnd.3 cells expressing *Prox1* exhibited discontinuous cell-cell junctions and an enlarged cell shape. **(C-D)** Three representative images of bEnd.3 cells expressing *Gfp* or *Prox1* labeled with p-MLC2 (cyan or grey) and F-Actin (magenta or grey) together with GFP (yellow) or PROX1 (yellow). Both F-Actin and p-MLC2 are downregulated in bEnd.3 cells expressing *Prox1* in comparison to bEnd.3 cells expressing *Gfp.* (E-G) Three representative images of bEnd.3 cells expressing *Gfp* or *Prox1* labeled with ZO-1 (cyan or grey) and CLDN5 (magenta or grey) together with GFP (yellow) or PROX1 (yellow). Boxed regions are magnified in the lower panels. bEnd.3 cells expressing Prox1 exhibited a significant reduction of CLDN5 expression. (G) Quantification of the CLDN5 fluorescence intensity in the TJ region Data are shown as mean ±SEM. n=20 fields of view from **4** independent experiments, *’**p<0.0001, as determined by unpaired I-test. Scale bars: 20 µmin (A-F).

The foregoing in vivo and in vitro studies demonstrate that *Prox1* expression in brain ECs leads to a decrease in the mRNA expression of *Cldn5* and a reduction of both junctional and cytoplasmic CLDN5 in brain ECs (Figure 7, E-F, Supplemental Figure 8A). Considering prior reports that suggest PROX1 functions as a transcriptional repressor in neural progenitors(64), hepatocytes(65), and cancers(66, 67), it is plausible that PROX1 regulates CLDN5 expression through direct transcriptional suppression of *Cldn5* gene. Analysis of a published whole-genome chromatin immunoprecipitation sequencing (ChIP-seq) using an anti-PROX1 antibody in human umbilical vein ECs (HUVECs) expressing *Prox1* reveals the presence of PROX1-binding sites at the promoter of *Cldn5* gene(68) (Supplemental Figure 8A). Likewise, we demonstrate that the endothelial *Prox1* expression leads to a decrease in the mRNA expression of *Ctnnb1/ß-catenin* and *Cd93,* and the whole-genome Prox1 ChIP-seq reveals the presence of PROX1-binding site at the *Ctnnb1/ß-catenin* and *Cd93* promoters. These findings imply that PROX1 may inhibit the promoter of *Cldn5*, *Ctnnb1*, or *Cd93* gene in brain ECs. Conversely, although the mRNA expression of *Mfsd2a* is also downregulated in brain ECs of *Prox1^iEC-OE^* mutant mice, a PROX1-binding site was not identified in its promoter region (Supplemental Figure 8A). Considering that the expression of *Mfsd2a* is known to be transcriptionally regulated by Wnt/ß-catenin signaling, the decreased expression of *Mfsd2a* in the mutants might be attributed to reduced ß-catenin level. Taken together, these data suggest that *Prox1* expression in brain ECs disrupts barrier integrity by reducing the expression of BBB-associated genes and Wnt/ß-catenin signaling in brain ECs (Supplemental Figure 8B).

## DISCUSSION

The immune privilege environment of the CNS parenchyma is maintained by unique immunological barriers, including the presence of the BBB and the absence of lymphatic vasculature. However, under pathological conditions such as brain tumors and AVMs, which compromise vascular integrity, there is an upregulation of LEC markers, including the LEC master regulator PROX1 and the vascular permeability marker PLVAP. Several lines of evidence suggest that PROX1 impairs BBB integrity by negatively regulating the expression of BBB-associated genes and Wnt/ß-catenin signaling. First, endothelial *Prox1* expression induces a hybrid blood-lymphatic phenotype in the developing CNS vasculature when activated during primitive BBB formation at embryonic stages, whereas it does not induce this phenotype during the BBB maturation at postnatal stages. Second, while *Prox1* is insufficient to induce conventional lymphatic vascular formation within the CNS parenchyma, it disrupts BBB integrity by downregulating TJ proteins and increasing transcytosis. These findings highlight the inhibitory effects of PROX1 on BBB development and maintenance. Third, PROX1 negatively regulates the expression of BBB-associated genes and Wnt/ß-catenin signaling in ECs.

Embryonic *Prox1* induction triggers the transformation of blood vessels into hybrid blood-lymphatic vessels, rather than the formation of conventional lymphatic vessels, within the brain parenchyma. Like Schlemm’s canal ECs in the eyes, *Prox1^iEC-OE^* mutant ECs express the LEC markers such as VEGFR3 and ITGα9, but not LYVE1 or PDPN, as well as the BEC markers such as PECAM1, ERG, and EMCN. Interestingly, the induction of *Prox1* postnatally does not result in a hybrid blood-lymphatic phenotype within the brain parenchyma, as *Prox1^iEC-OE^* mutant ECs fail to upregulate the expression of VEGFR3 and ITGα9. Given that the VEGF-C/VEGFR3 signaling is crucial for the early development of Schlemm’s canal ECs(42, 43), the lower VEGFR3 expression level may be insufficient to induce a hybrid blood-lymphatic phenotype.

Because *Vegfr3* is a direct target gene of PROX1(69), it is apparent that the postnatal CNS parenchyma establishes a microenvironment that prevents the upregulation of VEGFR3 expression in brain ECs. Detailed molecular mechanisms responsible for the suppression of LEC markers, such as VEGFR3, remain to be elucidated.

Despite the upregulation of LEC markers, including LYVE1, in ECs in brain tumors and AVMs, *Prox1^iEC-OE^* mutant ECs fail to differentiate into conventional LECs. These phenotypic differences suggest that the pathological microenvironments may provide additional signals that could induce the expression of the conventional LEC markers within the CNS parenchyma. The specific signals that alter the CNS non-permissive microenvironment for the development and growth of lymphatic vasculature are currently under investigation. Additionally, we should not discount the potential contribution of leptomeningeal LECs to the pathological brain parenchyma. While no report currently supports the invasion of leptomeningeal LECs into the brain parenchyma in mammals, a transient invasion of non-lumenized LECs into the injured brain parenchyma has been observed in a zebrafish model (70, 71).

Endothelial *Prox1* expression leads to an increased vascular leakage and BBB disruption when induced during both embryonic and postnatal stages. These data suggest PROX1’s inhibitory effects on barrier integrity. Indeed, the endothelial *Prox1* expression leads to decreased expression of TJ proteins such as CLDN5/Claudin-5 and ZO-1, along with the induction of PLVAP, a marker of high-permeability vasculature.

Although our TEM analysis does not reveal discontinuous cell-cell junctions or fenestrations in *Prox1^iEC-OE^* mutant capillaries, cultured bEnd.3 cells expressing *Prox1* displayed discontinuous cell-cell junctions. While the RhoA/ROCK signaling pathway typically induces the formation of radial actin stress fibers, increased contractility, and the disruption of cell-cell junctions(72, 73), this is not the case in the cultured bEnd.3 cells expressing *Prox1.* Instead, the discontinuous cell-cell junctions in these cultured bEnd.3 cells are likely the result of a combination of actin fiber destabilization and relaxation, along with decreased expression of TJ proteins. *Prox1^iEC-OE^* mutants in vivo do not exhibit discontinuous cell-cell junctions or fenestrations in brain capillaries, probably due to pericyte coverage. In addition, our findings indicate that *Prox1* expression leads to the upregulation of transcytosis, as indicated by reduced expression of *Mfsd2a,* a lipid transporter that limits transcytosis in the BBB, and elevated expression of *caveolin-1/CAV1*, accompanied by an increased number of endothelial vesicles. Given that *Mfsd2a* expression is transcriptionally regulated by Wnt/ß-catenin signaling in both in vivo(50, 51, 57, 61) and cultured bEnd.3 cells(51), PROX1 indirectly upregulates transcytosis by downregulating Wnt/ß-catenin signaling. Overall, these findings suggest that the endothelial *Prox1* expression leads to increased paracellular and intercellular permeability of the BBB.

Considering prior research indicating that impaired EC ß-catenin signaling results in increased paracellular and intercellular permeability of the BBB(46, 57–60), *Prox1* expression impacts ß-catenin signaling, as evidenced by reduced expression of *Ctnnb1/ß-catenin* and several effector and target genes including *Cldn5* and *Mfsd2a*. Likewise, PROX1 appears to inhibit the promoter of *Ctnnb1/ß-catenin* or *Cldn5* gene in ECs. How does PROX1 function as a transcriptional repressor in brain ECs? In hepatocytes, PROX1 interacts with the class I histone deacetylase HDAC3 to cooperatively repress gene transcription critical for maintaining lipid homeostasis(65). In colorectal cancer cells, PROX1 interacts with HDAC1 in the nucleosome remodeling and deacetylase (NuRD) complex to suppress Notch pathway(66). Indeed, HDAC2 mediates transcriptional regulation of BBB genes during BBB formation and maintenance(74). Thus, it is plausible that PROX1 may interact with the class I histone deacetylases such as HDAC2 to suppress the expression of *Ctnnb1/ß-catenin* or *Cldn5* in brain ECs.

Our studies clearly demonstrate that while CNS establishes a non-permissive microenvironment for the development and growth of conventional lymphatic vasculature under physiological conditions, endothelial *Prox1* expression leads to increased paracellular and intercellular permeability of the BBB. Despite the upregulation of LEC markers and BBB disruption observed in CNS ECs in brain tumors and AVMs, our genetic mouse models demonstrate that *Prox1* upregulation alone is sufficient to trigger vascular malformations and BBB disruption in the CNS vasculature. These findings indicate that tightly suppressing *Prox1* expression in CNS ECs may be necessary to preserve BBB integrity and prevent lymphatic vasculature formation in the CNS parenchyma. There are examples from non-CNS organs where *Prox1* suppression is crucial for maintaining the segregation between blood and lymphatic vasculatures.

For instance, deficiency in *Folliculin (FLCN)*, a tumor suppressor gene responsible for Birt-Hogg-Dubé (BHD) syndrome, leads to endothelial *Prox1* expression in veins, causing improper connections between blood vessels and lymphatic vessels(38). In zebrafish, the vascularization of the anal fin involves the transdifferentiation of pre-existing lymphatic vessels into blood vessels, with *Sox17* playing a crucial role in suppressing *Prox1* expression to facilitate the LEC-to-BEC transition(75).

Further studies are needed to elucidate the fundamental mechanisms underlying *Prox1* suppression in brain ECs and the absence of lymphatic vessels within the CNS parenchyma. In pathological conditions, dysregulation of *Prox1* expression could lead to BBB alterations. Understanding the molecular links between *Prox1* regulation and barrier disruption in disease states could facilitate the development of innovative therapeutic strategies, either to enhance drug delivery to the brain or to restore BBB function in the context of disease.

## Methods

### Sex as a biological variable

In this study, sex was not considered as a biological variable in embryos and neonates.

### Mice

The following mice (*Mus musculus*) were used in this study: C57BL/6J mice (The Jackson Laboratory), CD-1 mice (Charles River Laboratory), *Cadh5-BAC-Cre^ERT2^* mice(39), and *Prox1-GFP BAC* mice(36). *Rosa26-LSL-Prox1* mice have been generated in the Mukouyama Lab and the NHLBI Transgenic Core. For timed mating, the morning of the vaginal plug was considered E0.5. The *Cre*-mediated excision was induced by administering 1.5-3 mg tamoxifen (Sigma-Aldrich) by intraperitoneal injection (I.P.) at embryonic day (E)13.5, and embryos were harvested at E16.5 for analysis. For postnatal analysis, tamoxifen injection was performed I.P. (0.5 mg) to each pup at postnatal day (P)7 and analysis was performed at P10.

### Generation of *R26-LSL-Prox1* mice

The generation of *Rosa26-LSL-Prox1* mice was previously described(38). Briefly, a mouse *Prox1* coding sequence with 5’ FLAG-tag was knocked into the mouse *Rosa26 locus* using the CRISPR/Cas9 method in the NHLBI Transgenic Core. The *R26-LoxP-STOP-LoxP-Prox1* construct was co-microinjected along with *Cas9* mRNA and sgRNA into the pronuclei of fertilized mouse eggs. After culturing the injected embryos overnight, embryos that had reached the 2-cell stage of development were implanted into the oviducts of pseudopregnant foster mothers.

### scRNA-seq analysis of publicly available datasets

To evaluate lymphatic marker gene expressions, publicly available scRNA-seq datasets were utilized. For glioblastoma, three datasets were retrieved from the Gene Expression Omnibus (GEO) database under accession numbers GSE162631, GSE173278, and GSE184357. For brain metastasis, dataset was downloaded at: https://joycelab.shinyapps.io/braintime/. For AVMs, dataset was downloaded at: https://adult-brain-vasc.cells.ucsc.edu. For the glioblastoma datasets, the endothelial cell population was first subset from each dataset using R package Seurat. The three endothelial datasets were then integrated using the integration method provided by the Seurat package (76). Following integration, principal component analysis was performed for dimensional reduction. Uniform Manifold Approximation and Projection (UMAP) was then applied (dims = 1:30). For all three disease datasets (glioblastoma, brain metastasis, and AVMs), UMAP plots, violin plots, and dot plots were visualized using either scCustomize package in R (77) or Scanpy package in Python. To calculate average gene expressions of lymphatic markers (*PROX1*, *LYVE1*, and *FLT4*) and *PLVAP*, the AverageExpression function in Seurat was used.

### Histology and Immunofluorescence

Embryos and neonates harvested from timed pregnancies (morning of plug designated E0.5) were collected and washed in PBS and then fixed in 4% paraformaldehyde (PFA) overnight at 4°C with rotation. After washing in PBS, the fixed embryos and neonates were equilibrated in a 15% to 30% sucrose gradient at 4°C overnight. The tissues were embedded in Tissue-Tek O.C.T. Compound (Sakura). Cryosections were washed in PBS, permeabilized with PBS-T (0.5% Triton X-100) for 5-10 minutes when needed, and then blocked with 10% goat serum in PBS with 0.1% Triton X-100 or 1% bovine serum albumin with 0.1% Triton X-100 for 1-2 hours at room temperature. Primary antibodies with dilution 1:100-1:200 were incubated in blocking buffer at 4°C overnight. Fluorescence-conjugated secondary antibodies were used at the dilution of 1:300-1:500 in the blocking buffer, and sections were incubated with secondary antibodies for 1 hour at room temperature. After washing in PBS, sections were mounted with ProLong glass antifade mounting media (Thermo Fisher). Samples processed in similar manner, excluding the use of primary antibodies, were employed as negative controls to verify the staining’s specificity in sections.

### Tissue clearing, whole-mount immunostaining, and confocal imaging

CUBIC method was used for tissue clearing as previously described(78). Briefly, after washing the fixed tissues in PBS, the tissues were incubated with CUBIC reagent-1 (25 wt% urea, 25 wt% *N,N,N’,N’*-tetrakis(2-hydroxypropyl) ethylenediamine, and 15% (v/v) Triton X-100) for 1-2 days at room temperature with rotation. The tissues were blocked with 10% goat serum in PBS with 0.1% Triton X-100 or 1% bovine serum albumin with 0.1% Triton X-100 for 12-24 hours. Primary (1:300) and secondary antibodies (1:500) were diluted in the blocking buffer and all washes were performed in PBS-T (0.05% Triton X-100 in PBS) with rotation. After whole-mount immunostaining, the tissues were balanced with sucrose (20%) and incubation was performed with CUBIC reagent-2 (50 wt% sucrose, 25 wt% urea, 10 wt% 2,2′,2′’-nitrilotriethanol, and 0.1% (v/v) Triton X-100) at room temperature with rotation in the dark for 1 day. Cubic reagent-2 was used as mounting medium for the confocal acquisition. All confocal microscopy was carried out on a Leica TCS SP5 microscope. Optical z-stack projections were generated with FIJI or Imaris software using a maximal intensity algorithm.

### Flow cytometry

Embryonic or postnatal brains were isolated in cold HBSS medium (Thermo Fisher). The brain tissues were minced into small pieces and digestion solution (0.05% DNase I, 0.1% collagenase, 0.3% dispase, in Leibovitz’s L-15 medium (Thermo Fisher) was incubated at 37°C for 45 minutes with agitation every 5-10 minutes. After dissociation, remaining clumps of cells were filtered through 70-μm filters and washed with cold FACS buffer (1% BSA, 0.1 M HEPES, 1x Pen-Strep, 0.025% DNase I, in L15 medium). Cells were centrifuged and resuspended in cold FACs buffer. Negative selection with magnet beads was performed to eliminate erythrocytes and myeloid cells. Briefly, cells were incubated with mouse anti-Ter119 (eBioscience, 1:100), mouse anti-CD45 (Biolegend, 1:100) for 30 minutes on ice. After washing with FACS buffer, cells were incubated with anti-rat IgG conjugated magnetic beads for 30 minutes. Negative selection was performed using a magnetic stand with 2 minutes of incubation per sample. Final samples were stained with the following antibody mix. All flow cytometry analyses were done on BD LSR Fortessa equipped with Diva Software. Cell sorting was performed using BD FACSMelody, BD FACSymphony or BD FACSAria Fusion.

Unstained samples, single-color staining, and fluorescence minus one (FMO) were used to establish the proper compensation and gating. In all samples, debris, blood cells and myeloid cells were gated out by DAPI, Ter119, and CD45 staining. Antibodies used for cytometry are listed: rat monoclonal anti-Ter119-BV785 (Biolegend, 1;100), rat monoclonal anti-CD45-BV785 (Biolegend, 1:100), rat monoclonal anti-CD31-PECy7 (eBioscience, 1:100), rat monoclonal anti-CD140b-APC (eBioscience, 1:100), rat monoclonal anti-NG2 AF488 (Millipore sigma, 1:100), rat monoclonal anti-LYVE-1-PE (MBL, 1:100). Data was analyzed using FlowJo software (BD biosciences).

### Quantitative real time PCR

mRNA was extracted from embryonic, postnatal brain, and skin ECs using PicoPure RNA Isolation Kit (Thermo Fisher), according to the manufacturer’s instructions. The mRNA was converted to cDNA using SuperScript III Reverse Transcriptase (Thermo Fisher). Quantitative real time (qRT)-PCR was performed in triplicate with Power SYBR™ Green Master Mix 2x (Roche). Relative quantification of each transcript was obtained by normalizing against *GADPH* transcript abundance. The general cycling conditions were as follows: one initial hold for 3 minutes at 95°C, followed by 40x cycles of 10-sec denaturation (95°C) and 45 seconds of annealing/extension at 60°C. The sequences of oligonucleotides for qRT-PCR are listed in Supplementary Methods:

### Blood-brain barrier permeability assay

Embryos were harvested at E16.5. Once the placenta and yolk sac were removed, 3kDa Dextran Texas-red (Invitrogen) was injected into the left ventricle of the heart (10µg in PBS) using a mouth pipette and glass capillaries. Injected embryos were incubated in HBSS medium for 5 min at room temperature, followed by fixation in 10% PFA/PBS for 2 hours at room temperature and rotation. After fixation, the embryos were washed three times with PBS, and then were equilibrated in a 15% to 30% sucrose gradient at 4°C overnight. The following day, tissues were embedded in Tissue-Tek O.C.T. Compound (Sakura) for cryosections or tissue-clearing and whole-mount immunostaining. Those embryos where the heart was not pumping correctly were not considered for analysis.

Neonates were harvested at P10 and injected intraperitoneally with 250µg 3kDa Dextran Texas-Red (Invitrogen) per 20g mouse or 100µg 1kDa Cadaverine (Thermo Fisher) per 20g mouse, as previously reported(79). After 2 hours, pups were euthanized, and brain tissues were harvested for fixation and posterior analysis. Leakage was determined by making a mask of the vasculature area using the PECAM1 channel, then assessing the dextran or cadaverine signal outside of the vasculature.

### Cell culture

Commercially available bEnd.3 cells (ATCC) were cultured in DMEM complete (ATCC) supplemented with 10% FBS, and 10 mM Penicillin/Streptomycin. Commercially available primary RBMVECs (Cell applications) were cultured in rat brain endothelial cell growth medium (Sigma) supplemented with 10 mM Penicillin/Streptomycin, according to the manufacturer’s instructions. All cells were maintained at 37°C and 5% CO_2_.

For lentivirus infection, cells were seeded onto 12-well glass chamber slides (Ibidi) coated with 10 µg/ml fibronectin (Millipore-Sigma) or attachment factor solution (Cell application). Once cells reached ∼60-70% confluency, cell medium was removed and fresh cell medium containing 1 mg/ml polybrene (Vector builder) and lentivirus expressing *Gfp* or *Prox1* was added (MOI 5-10). For each experiment, three separate plates were seeded with cells, one with no lentivirus, one with *Gfp*-lentivirus, and one with *Prox1*-lentivirus. Two days after the lentivirus infection, the cell medium was changed, and cells were fixed with 4% PFA for 10 minutes at RT when reached a confluent monolayer. Immunostaining was performed as described above. Cells were permeabilized with 0.1% Triton X-100 for 10 minutes at RT followed by blocking with BSA buffer for 1-2 hours at room temperature. Primary antibodies were incubated overnight at 4°C and secondary antibodies (1:400) the following day for 1-2 hours at room temperature. After the washing steps, Ibidi chambers were removed, and slides were mounted normally using ProLong mounting media (Thermo Fisher).

The following primary antibodies were used in cell culture: Goat anti-Prox1 (R&D, 1:100), mouse anti-Claudin-5 (Invitrogen, 1:100), rabbit anti-ZO-1 (Proteintech, 1:300), goat anti-GFP (Abcam 1:200), rabbit anti-pMLC2 (Cell signaling, 1:200), and Alexa Fluor 568 Phalloidin (Invitrogen, 1:500). RBMVECs at passages 3-5 were used for experiments. All confocal microscopy was carried out on a Leica TCS SP5 microscope using a 63x oil objective.

### Transmission Electron Microscopy

Postnatal brains were harvested and fixed by immersion in a 0.1M sodium cacodylate-buffered mixture (2.5% glutaraldehyde and 4% PFA) for 2 hours at RT followed by overnight incubation in 4% PFA at 4°C. The next day, tissues were washed two times in 0.1M sodium cacodylate buffer and then cut in 200 μm-thick free-floating sections using a vibratome. Sections were then post-fixed in 2% osmium tetroxide and 1.5% potassium ferrocyanide and stained overnight in 1% UA. The following day samples were dehydrated in graded ethanol series and infiltrated with resin (Embed-812) and baked at 60°C for 48 hours. Ultrathin sections (65-70 nm) were cut on an ultramicrotome (Leica EM UC7), and digital micrographs were acquired with a JOEL JEM 1200 EXII (80 kV) equipped with an AMT XR-60 digital camera.

### Quantification and Statistical Analysis

All data were collected from at least three independent experiments as indicated. The actual number of independent biological replicates are specified, wherever applicable, in the relevant figure legends. Statistical analyses were performed using Prism (GraphPad v9.0). Data are presented as mean values ±SEM. The Shapiro-Wilk test was used to check the normality of data distribution. When the normality assumption was met, unpaired t-test was applied to assess the significance. For all the images included across the manuscript, the most representative examples reflecting the typical phenotype were selected.

### Study approval

All animal procedures were approved by the National Heart, Lung, and Blood Institute (NHLBI) Animal Care and Use Committee in accordance with NIH research guidelines for the care and use of laboratory animals.

### Data availability

All data in the manuscript is included in the Support Data Values file.

## ACKNOWLEGMENTS

Thanks to P. Dagur, M. Lopez-Ocasio, and K. Keyvanfar of the NHLBI Flow Cytometry Core for FACS assistance; Z. Syed of the NHLBI Electron Microscopy Core for TEM assistance; T. Markowitz and N. Redekar of Research Technology Branch in the National Institute of Allergy and Infectious Diseases (NIAID) for ChIP-seq data analysis; T. Clark and the staff of NIH Bldg50 animal facility for assistance with mouse breeding and care. Thanks also to C. Conrad of the Developmental Therapeutics and Pharmacology Unit in National Institute of Neurological Disorders and Stroke (NINDS) for sharing reagents; C. Waterman, R. Fischer, A. Pasapera in the Laboratory of Cell and Tissue Morphodynamics (NHLBI), and V. Bautch and D. Buglak (University of North Carolina) for sharing reagents and valuable discussion. Thanks also to K. Gill for laboratory management and technical support, V. Sam for administrative assistance, and members of the Laboratory of Stem Cell and Neuro-Vascular Biology for technical help and thoughtful discussion. S. González-Hernández is supported by the NHLBI Lenfant Biomedical Fellowship. This work was also supported by the Intramural Research Program of the NHLBI, NIH (HL006115-14 to Y-S Mukouyama).

## Author contributions

S. G-H conducted all the experiments, and also contributed to the conceptualization, writing and editing of the manuscript. Y. S., C. L., W. L., and C. L were responsible for generating and conducting the primary characterization of *R26-LSL-Prox1* mice. S. J. provided valuable reagents. Y. K. provided *Cdh5-Cre^ERT2^* mice. Y-S. M. contributed through project supervision, discussion, and writing and editing of the manuscript.

