## Supplemental information for "ENDOTHELIAL PROX1 INDUCES BLOOD-BRAIN BARRIER DISRUPTION IN THE CENTRAL NERVOUS SYSTEM"

**Supplementary Figures**

Supplemental Figure 1, related to Figure 1: Expression of LEC markers and BBB-associated markers in endothelial cells within human brain tumors and vascular malformations.

Supplemental Figure 2, related to Figure 2: Lack of lymphatic vasculature in the brain parenchyma.

Supplemental Figure 3, related to Figure 3: Characterization of $Prox1^{iEC\text{-}OE}$ mutant and their WT control littermate embryos.

Supplemental Figure 4, related to Figure 4: Endothelial *Prox1* expression induces aberrant lymphatic vasculature in non-CNS tissues.

Supplemental Figure 5, related to Figure 5: Blood-brain barrier disruption in E16.5 $Prox1^{iEC\text{-}OE}$ mutant embryos.

Supplemental Figure 6, related to Figure 6: Postnatal induction of *Prox1* disrupts the matured blood-brain barrier without inducing a hybrid blood-lymphatic phenotype.

Supplemental Figure 7, related to Figure 7: *Prox1* overexpression in RBMVECs induces abnormal tight junctions.

Supplemental Figure 8, related to Figure 7: PROX1 negatively regulates the expression of *Ctnnb1* and BBB-related *Cldn5* and *Cd93.*

**Supplementary Methods**

Supplemental Table 1: List of oligonucleotides for qRT- PCR

### Supplemental Figure 1

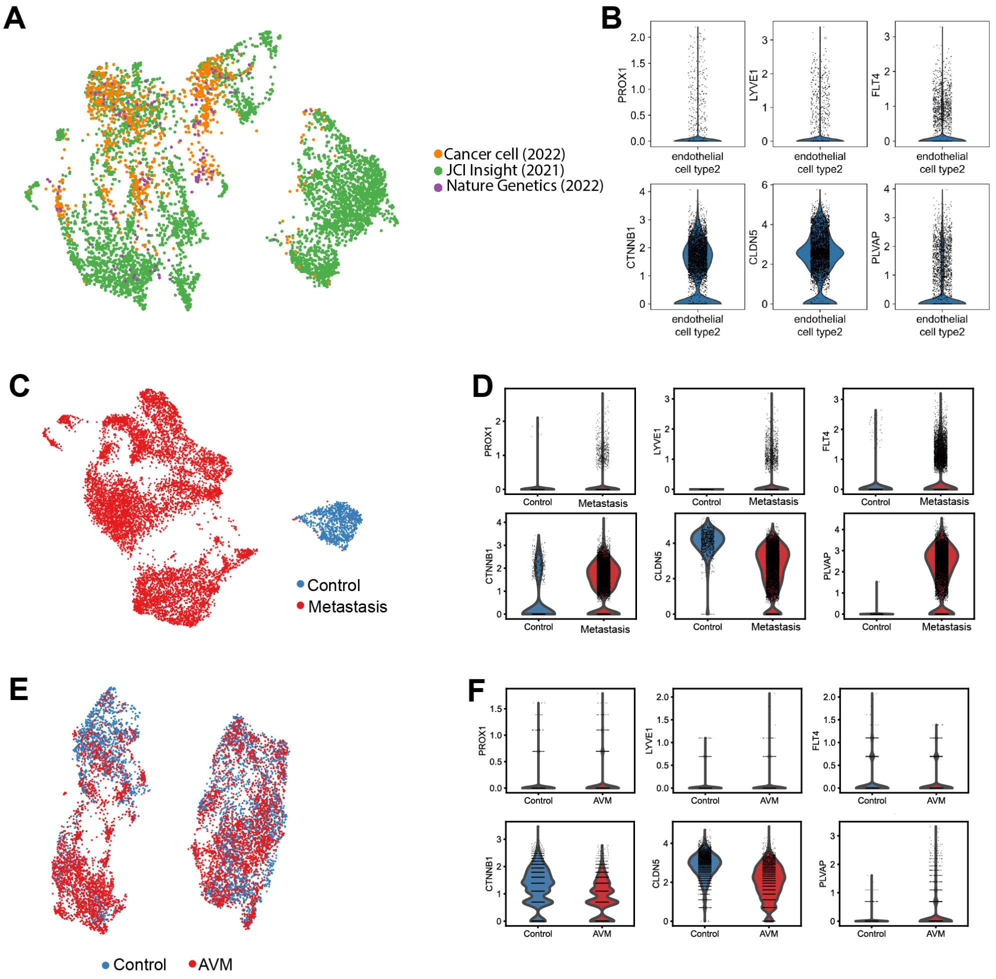


**Supplemental Figure 1. Expression of LEC markers and BBB-associated markers in endothelial cells within human brain tumors and vascular malformations.**
**(A-B)** UMAP (A) and violin plots (B) of scRNA-seq data from glioblastoma datasets display endothelial cell (EC) clusters expressing LEC markers (PROX1, LYVE1, FLT4), BBB-associated markers (CTNNB1, CLDN5), along with the vascular permeability marker PLVAP. **(C-D)** UMAP (C) and violin plots (D) of scRNA-seq data from control and metastatic brain tumor datasets display endothelial cell (EC) clusters expressing LEC markers (PROX1, LYVE1, FLT4), BBB-associated markers (CTNNB1, CLDN5), along with the vascular permeability marker PLVAP. **(E-F)** UMAP (E) and violin plots (F) of scRNA-seq data from control and AVM datasets display endothelial cell (EC) clusters expressing LEC markers (PROX1, LYVE1, FLT4), BBB-associated markers (CTNNB1, CLDN5), along with the vascular permeability marker PLVAP.

.

### Supplemental Figure 2

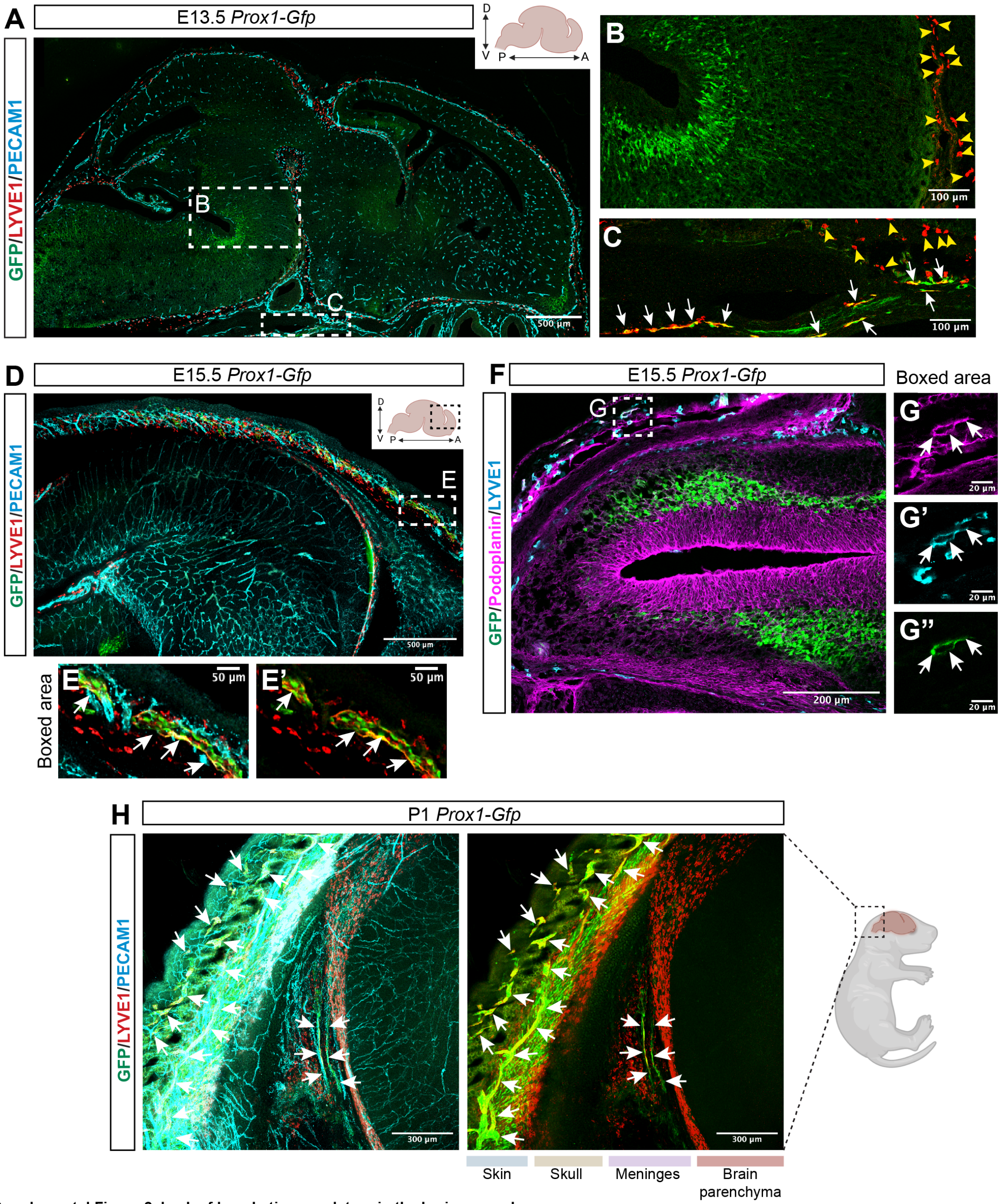


**Supplemental Figure 2. Lack of lymphatic vasculature in the brain parenchyma.**
**(A-C)** A sagittal view of E13.5 *Prox1-Gfp* head labeled with LYVE1 (red) and PECAM1 (cyan). The boxed regions in (A) are magnified in (B and C). Arrows in (C) indicate PECAM1+/LYVE1+/Prox1-GFP+ lymphatic vessels outside of the brain parenchyma, while yellow arrowheads in (B and C) indicate LYVE1+/PECAM1-/Prox1-GFP- macrophages. Scale bars: 500 µm in (A) and 100 µm in (B and C). **(D-G)** A sagittal view of E15.5 *Prox1-Gfp* head labeled with LYVE1 and PECAM1 (D-E) or Podoplanin and LYVE1 in (F-G). The boxed regions in (D and F) are magnified in (E and E') and (G-G"), respectively. Arrows indicate PECAM1+/LYVE1+/Prox1-GFP+ lymphatic vessels in (E and E') and Podoplanin+/LYVE1+/Prox1-GFP+ Lymphatic vessels in (G-G") in the skin surrounding the skull. Scale bars: 500 µm in (D), 200 µm in (F), 50 µm in (E and E'), and 20 µm in (G-G"). **(H)** A sagittal view of whole-mount immunostaining of P1 *Prox1-Gfp* head labeled with PECAM1 (cyan) and LYVE1 (red). Arrows indicate PECAM1+/LYVE1+/Prox1-GFP+ lymphatic vessels in the meninges and skin. Scale bars: 500 µm in (A and D), 30 µm in (I and J), and 10 µm in (H). Scale bars: 300 µm. The illustrations are created with BioRender.com.

### Supplemental Figure 3

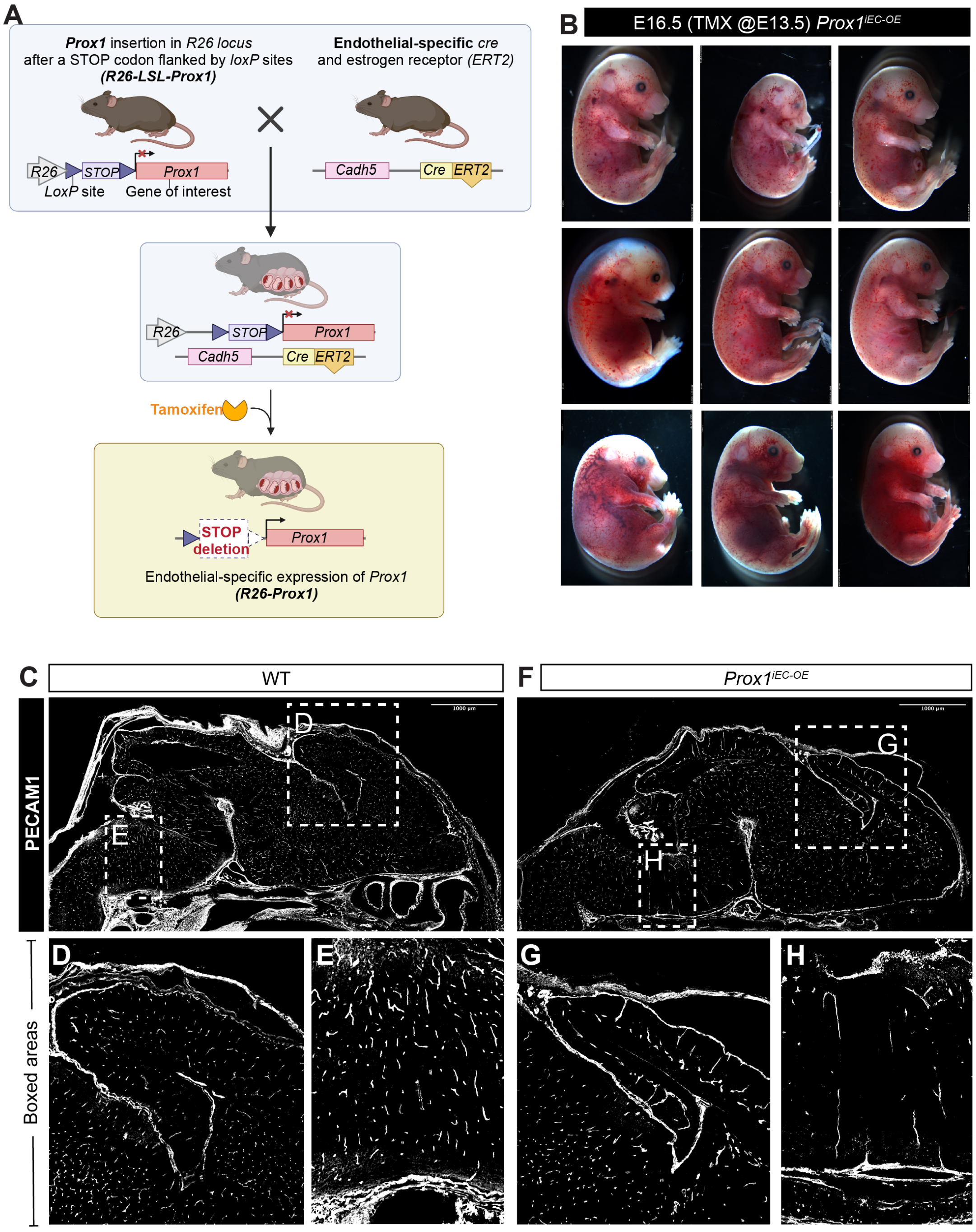


**Supplemental Figure 3: Characterization of Prox1iEC-OE mutant and their WT control littermate embryos.**
**(A)** Schematic diagram depicting the generation of *Prox1*$^{iEC-OE}$ mice and EC-specific *Prox1* expression. **(B)** Gross appearance of E16.5 *Prox1*$^{iEC-OE}$ mutant embryos when induced by tamoxifen at E13.5. *Prox1*$^{iEC-OE}$ mutant embryos show edemas, hemorrhagic manifestation, and blood-filled lymphatics in the skin, with variations in phenotypic severity. **(C-H)** A sagittal view of E16.5 *Prox1*$^{iEC-OE}$ mutant and their WT control littermate brains labeled with PECAM1 (grey). The boxed regions in (C and F) are magnified in (D-E and G-H), respectively. Scale bars: 1000 μm in (C and F). The illustrations are created with BioRender.com.

### Supplemental Figure 4

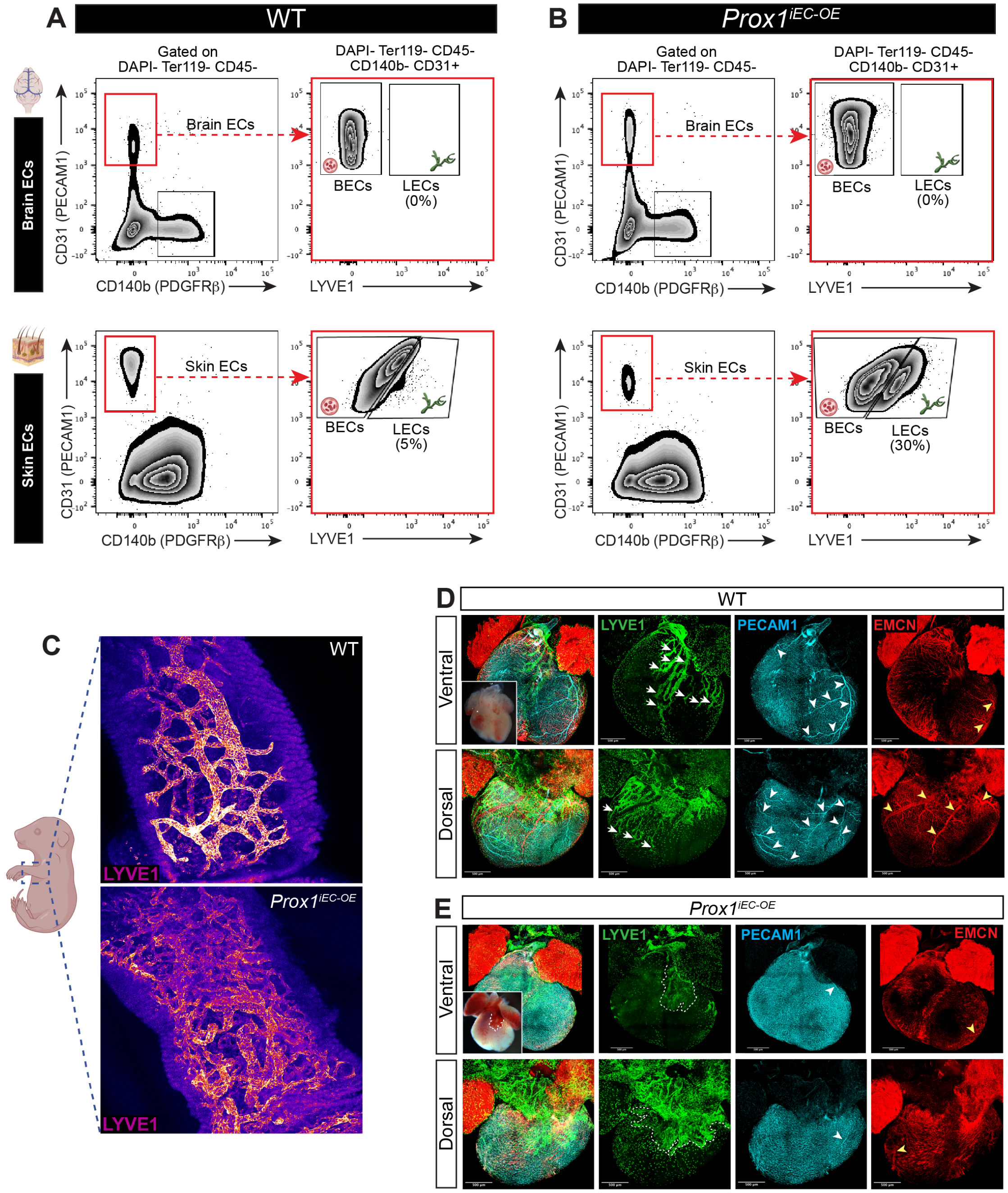


**Supplemental Figure 4. Endothelial *Prox1* expression induces aberrant lymphatic vasculature in non-CNS tissues.**
**(A-B)** Flow cytometry analysis of BECs (DAPI-/Ter119-/CD45-/CD140b-/CD31+/LYVE1-) and LECs (DAPI-/Ter119-/CD45-/CD140b-/CD31+/LYVE1+) in brain and skin from E16.5 *Prox1*$^{iEC\text{-}OE}$ mutant and their WT control littermate embryos. Ectopic expression of Prox1 increases in the proportion of LECs in the skin but not in the brain. **(C)** Whole-mount immunostaining of limb skin from E16.5 *Prox1*$^{iEC\text{-}OE}$ mutant and their WT control littermate embryos labeled with LYVE1 (fire channel). **(D-E)** Whole-mount immunostaining of heart ventricle from E16.5 *Prox1*$^{iEC\text{-}OE}$ mutant and their WT control littermate embryos labeled with LYVE1 (green), PECAM1 (cyan) and EMCN (red). Inset images indicate blood-filled cardiac lymphatic vasculature in the ventral surface of the mutant heart ventricle. Arrows indicate LYVE1+ lymphatic vasculature in the ventral and dorsal surface of the WT control heart ventricle. Dashed outlines indicate aberrant LYVE1+ lymphatic vasculature in the ventral and dorsal surface of the mutant heart ventricle. Arrowheads in the PECAM1 staining indicate large-diameter coronary arteries in the myocardium, while yellow arrowheads in the EMCN staining indicate large-diameter coronary veins in the subepicardium. Scale bars: 500 µm. The illustrations are created with BioRender.com.

### Supplemental Figure 5

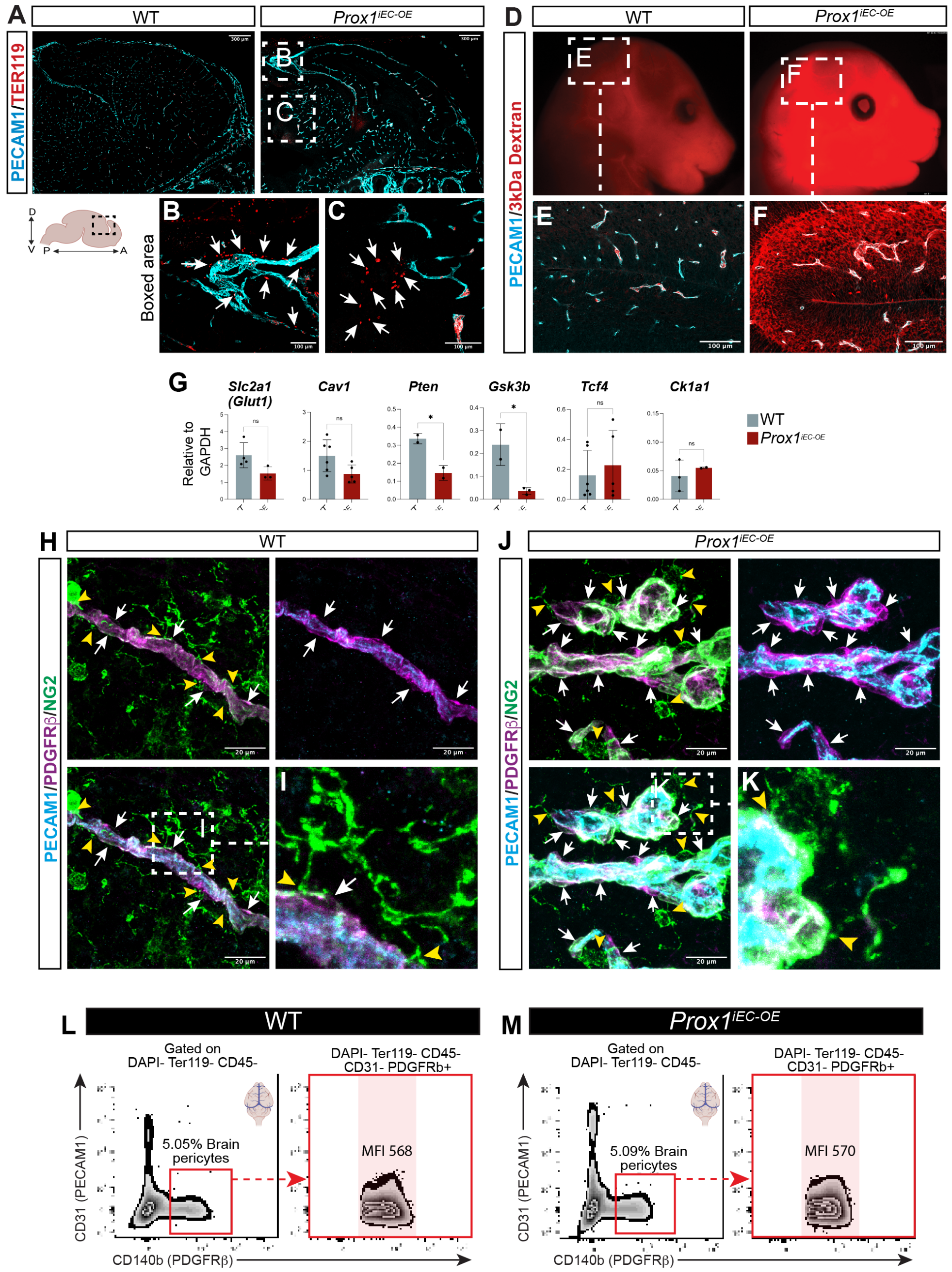


**Supplemental Figure 5. Blood-brain barrier disruption in E16.5** ***Prox1***$^{iEC-OE}$ **mutant embryos.**
**(A-C)** Section immunostaining of E16.5 *Prox1*$^{iEC-OE}$ mutant and their WT control littermate brains with PECAM1 (cyan) and Ter119 (red). The boxed regions in (A, Prox1iEC-OE mutants) are magnified in (B and C). Arrows in (B and C, *Prox1*$^{iEC-OE}$ mutants) indicate Ter119+ blood cell extravasation. Scale bars: 300 µm in (A) and 100 µm in (B and C). **(D-F)** A tracer leakage assay of E16.5 *Prox1*$^{iEC-OE}$ mutant and their WT control littermate brains with 3kDa Dextran Texas-Red. The upper panels indicate gross appearance of E16.5 *Prox1*$^{iEC-OE}$ mutant and their WT control littermate heads. The boxed regions in the upper panels largely correspond to the immunostaining images labeled with PECAM1 (cyan), displayed in the lower panels (E and F), The mutant brains exhibited extensive BBB leakage. Scale bars: 100 µm. **(G)** Relative mRNA expression levels of BBB-related genes such as *Slc2a1/Glut1*, *Cav1* and *Pten*, and *ß-catenin* target genes such as *Gsk3b*, *Tcf4*, and *Ck1a1* in FACS-isolated brain ECs from E16.5 *Prox1*$^{iEC-OE}$ mutant and their WT control littermate brain. Graphs show mean normalized expression ±SEM; n=2-6 biological samples obtained from FACS-isolated brain ECs from individual experiments. *$p<0.005$, as determined by unpaired t-test. **(H-K)** Section immunostaining of E16.5 *Prox1*$^{iEC-OE}$ mutant and their WT control littermate brains with PECAM1 (cyan), PDGFRß (magenta), and NG2 (green). The boxed regions in the lower left panels of WT control (H) and mutants (J) are magnified in (I and K), respectively. Arrows indicate PDGFRß+/NG2+ pericyte coverage of PECAM1+ capillaries. Yellow arrowheads indicate NG2+/PDGFRß- oligodendrocytes associating with capillaries. Scale bars: 20 µm. **(L-M)** Flow cytometry analysis of pericytes (DAPI-/Ter119-/CD45-/CD31-/CD140b+ in E16.5 *Prox1*$^{iEC-OE}$ mutant and their WT control littermate brain. The expression level of CD140b (PDGFRß) marker is the same, as determined by the maximal fluorescence intensity (MFI). The illustrations are created with BioRender.com.

### Supplemental Figure 6

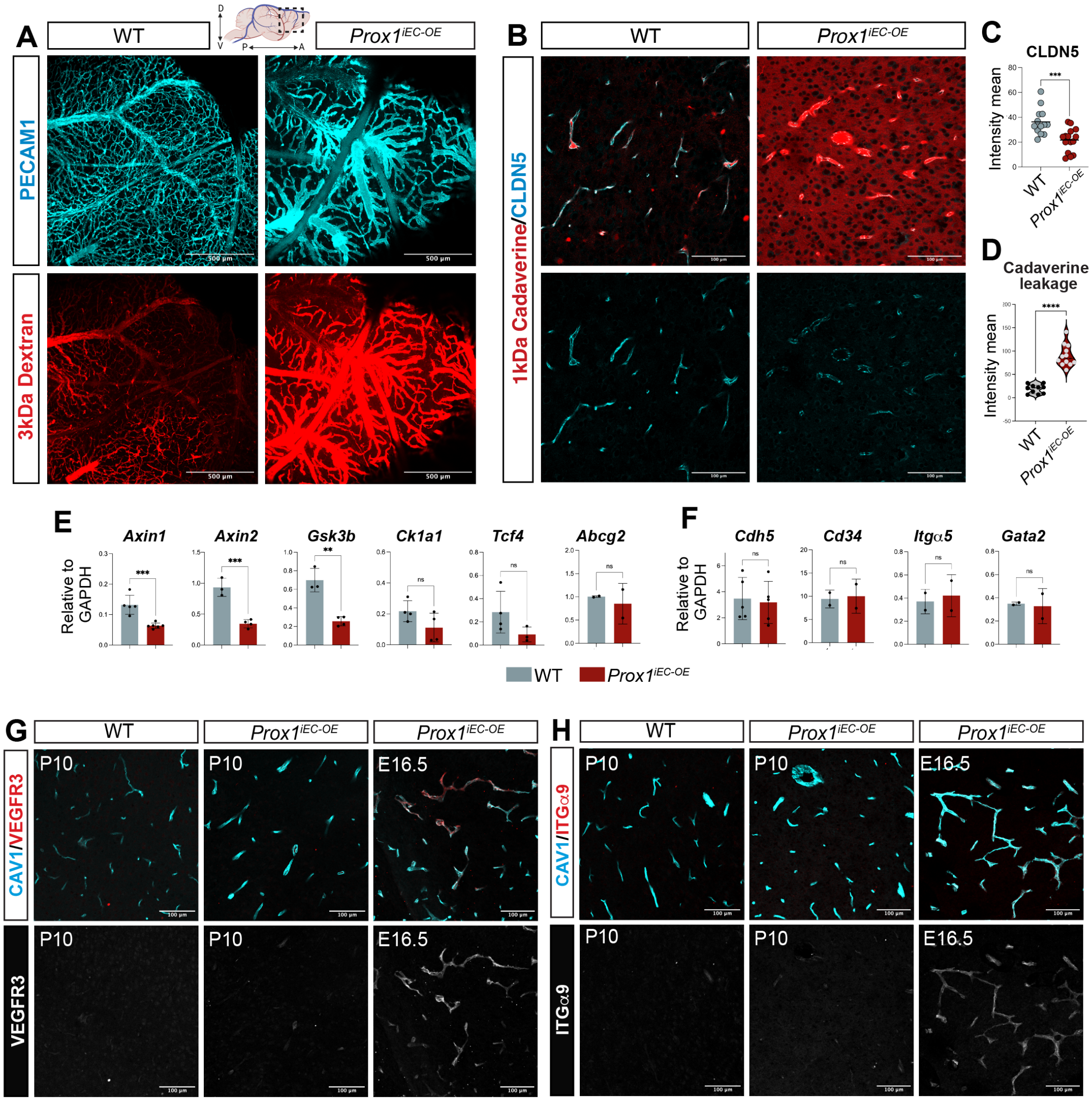


**Supplemental Figure 6. Postnatal induction of *Prox1* disrupts the matured blood-brain barrier without inducing a hybrid blood-lymphatic phenotype.**

**(A)** A surface view of whole-mount immunostaining of P10 *Prox1$^{iEC-OE}$* mutant and their WT control littermate brains with 3kDa Dextran Texas-Red tracer (red) and PECAM1 (cyan). Scale bars: 500 µm. **(B-D)** Section immunostaining of P10 *Prox1$^{iEC-OE}$* mutant and their WT control littermate brain with 1kDa Cadaverine tracer (red) and CLDN5 (cyan). (C-D) Quantifications of the fluorescence intensity mean of CLDN5 in (C) and cadaverine extravasation outside of the brain vasculature in (D) in *Prox1$^{iEC-OE}$* mutant and their WT control littermate brain. Dots corresponds to random fields of view from at least 3 different WT and mutant embryos. ***p<0.0005; ****p<0.0001, as determined by unpaired t-test. Scale bars: 100 µm. **(E-F)** Relative mRNA expression levels of ß-catenin target genes such as *Axin1, Axin2, Gsk3b, Ck1a1, Tcf4,* and *Abcg2* (E) and BEC markers such as *Cadh5, Cd34, Itgα5* and *Gata2* (F) in FACS-isolated brain ECs from P10 *Prox1$^{iEC-OE}$* mutant and their WT control littermate brain. Graphs show mean normalized expression ±SEM; n=2-4 experiments obtained from 4 individual FACS experiments. ** p<0.001 ***p<0.0005, as determined by unpaired t-test. **(G-H)** Section immunostaining of P10 and E16.5 *Prox1$^{iEC-OE}$* mutant and their WT control littermate brains with CAV1 (cyan) as a vascular marker, together with VEGFR3 (red and grey) in (G) or ITGα9 (red and grey) in (H). Postnatal *Prox1* induction does not upregulate VEGFR3 and ITGα9, unlike embryonic *Prox1* induction. Scale bars: 100 µm. The illustrations are created with BioRender.com.

### Supplemental Figure 7

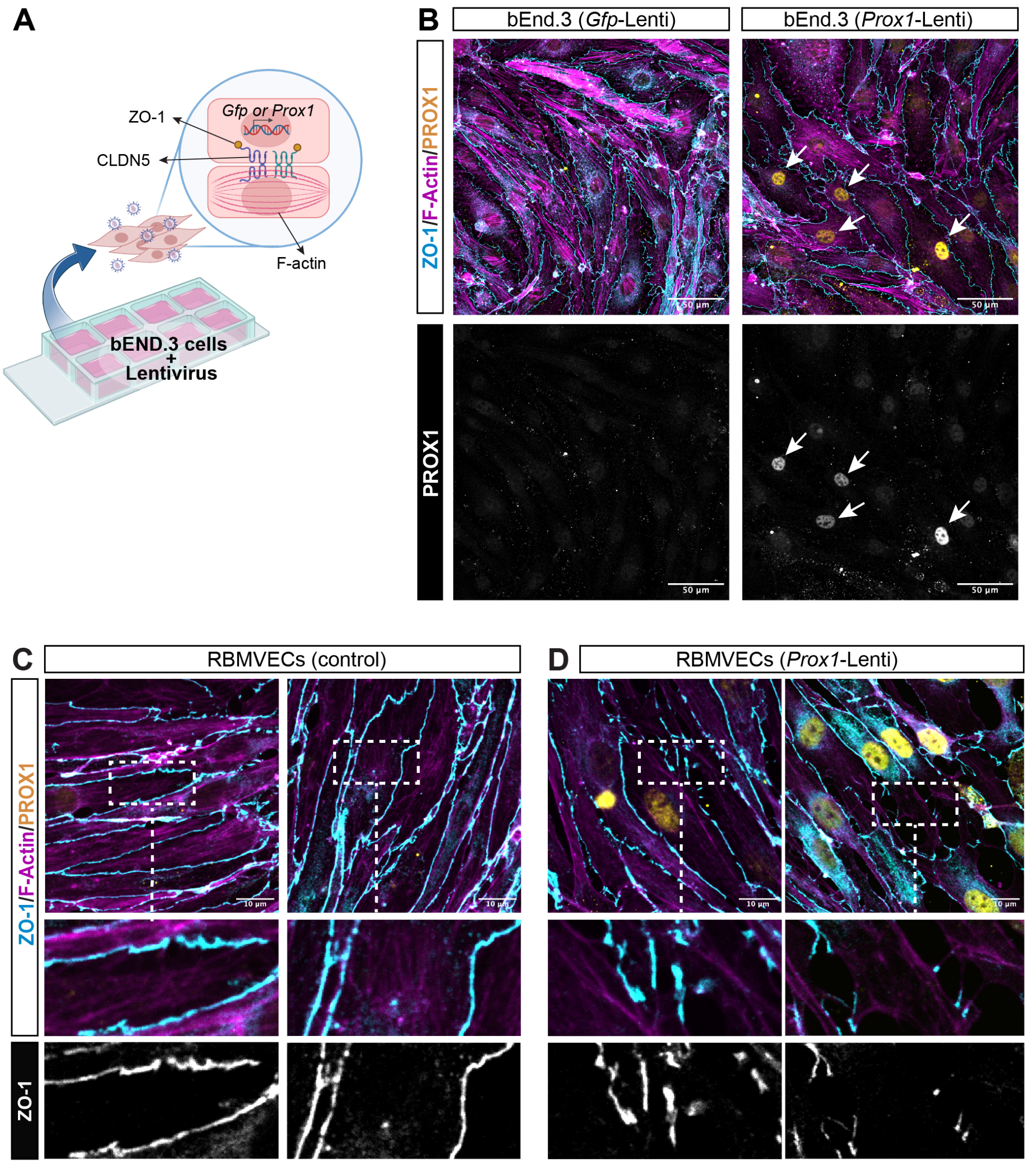


**Supplemental Figure 7. *Prox1* overexpression in RBMVECs induces abnormal tight junctions.**
**(A)** Schematic diagram depicting the preparation of bEnd.3 cells expressing *Gfp* or *Prox1*. bEnd.3 cells are cultured in 8-12 well Ibidi chamber slides and transduced with empty lentivirus expressing *Gfp* (Gfp-Lentivirus) or *Prox1* gene (Prox1-Lentivirus) to study the effect of *Prox1* expression in the tight junctions and actin filaments. **(B)** Immunostaining of bEnd.3 cells expressing *Gfp* or *Prox1* with PROX1 (yellow and grey), ZO-1 (cyan) and F-Actin (magenta). Arrows in bEnd.3 cells expressing *Prox1* indicate PROX1 expression. bEnd.3 cells expressing *Gfp* do not express PROX1. Scale bars: 50 µm. **(C-D)** Two representative images of cultured RBMVEC rat brain ECs with ZO-1 (cyan and grey), F-actin (magenta) and PROX1 (yellow). The boxed regions in the upper panels are magnified in the lower panels. RBMVECs expressing *Prox1* exhibited discontinued cell-cell junctions. Scale bars: 10 µm. The illustrations are created with BioRender.com.

### Supplemental Figure 8

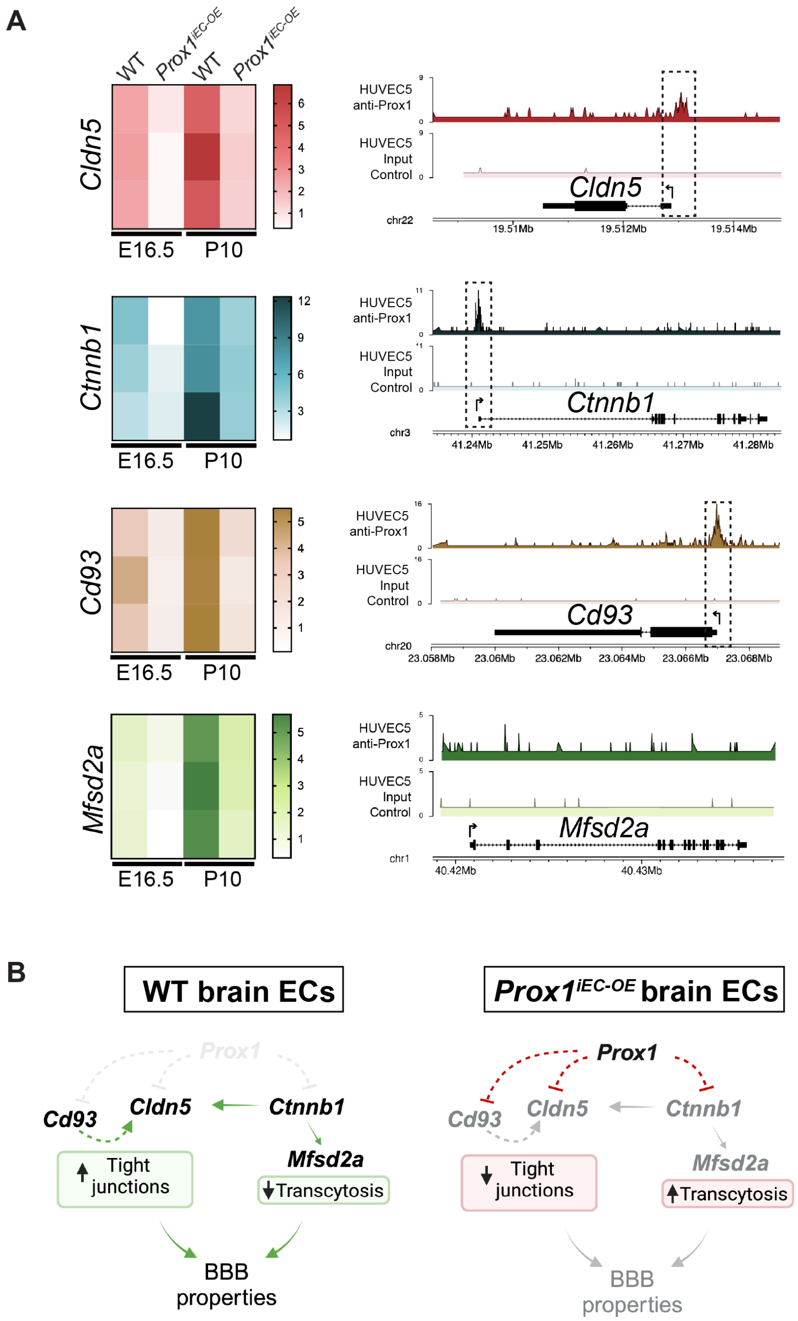


**Supplemental Figure 8. PROX1 negatively regulates the expression of *Ctnnb1* and BBB-related *Cldn5* and *Cd93*.**
**(A)** Heatmaps show the downregulation of *Cldn5, Ctnnb1, Cd93* and *Mfsd2a* expression in E16.5 or P10 *Prox1*$^{iEC\text{-}OE}$ brain ECs in comparison to their WT control littermates. A published whole-genome ChIP-seq study using an anti-PROX1 antibody in human umbilical vein ECs (HUVECs) expressing *Prox1* (Wong et al.68) reveals the presence of PROX1-binding sites at the promoter of *Cldn5, Ctnnb1*, and *Cd93*, but not *Mfsd2a*. This data is available online through the Gene Expression Omnibus (GEO) under reference number GSE71230. **(B)** Schematic illustrating the potential molecular mechanism by which the ectopic expression of *Prox1* induces BBB breakdown. The ectopic expression of Prox1 in brain ECs decreases the expression of *Cldn5, Ctnnb1* and *Cd93*, all critical for developing and maintaining the BBB properties. PROX1 directly downregulates *Cldn5* expression. Moreover, reduced expression of *Ctnnb1* and *Cd93* also contributes to deficient *Cldn5* expression, therefore affecting to tight junctions. Additionally, reduced *Ctnnb1* expression impairs the expression of *ß-catenin/Ctnnb1* sensitive *Mfsd2a*, inducing to abnormal transcytosis activity.

**Supplementary Methods:**

**Supplemental Table 1:** List of oligonucleotides for qRT- PCR

GAPDH Fw: CTGCACCACCAACTGCTTAG, GAPDH Rev: TCTCATCATACTTGGCAGGT,

Prox1 Fw: AGAAGGGTTGACATTGGAGTGA, Prox1 Rev: TGCGTGTTGCACCACAGAATA,

Claudin-5 Fw: ACATGCAGTGCAAGGTGTAT, Claudin-5 Rev: GGTAACAAAGAGTGCCACCA,

Plvap Fw: GCTGGTACTACCTGCGCTATT, Plvap Rev: CCTGTGAGGCAGATAGTCCA,

Sox17 Fw: ACGCTAGCTCAGCGGTCTACTATT, Sox17 Rev: AGGGATTTCCTTAGCGCTTCCAGG,

Ctnnb1 Fw: GTTCGCCTTCATTATGGACTGCC, Ctnnb1 Rev: ATAGCACCCTGTTCCCGCAAAG,

Lef1 Fw: ACTGTCAGGCGACACTTCCATG, Lef1 Rev: ACTGTCAGGCGACACTTCCATG,

GLUT1 Fw: GCTTCTCCAACTGGACCTCAAAC, GLUT1 Rev: ACGAGGAGCACCGTGAAGATGA,

Axin1 Fw: ACGAGGAGCACCGTGAAGATGA, Axin1 Rev: GCCCATTGACTTGGATACTCTCC,

Axin2 Fw: ATGGAGTCCCTCCTTACCGCAT, Axin2 Rev: GTTCCACAGGCGTCATCTCCTT,

FZD4 Fw: ACTTTCACGCCGCTCATCCAGT, FZD4 Rev: TCTCAGGACTGGTTCACAGCGT,

Fgfbp1 Fw: ACGGAGCCAAACAGGGTCAAAG, Fgfbp1 Rev: GGTCTTCCTCTGGTTGAGCACA,

APCDD1 Fw: CGGTGTGCTCTCATCTAAGGTC, APCDD1 Rev: CCCACTGAAGACATTGAGGAGG,

ITGa5 Fw: CTTCTCCGTGGAGTTTTACCG, ITGa5 Rev: CTTCTCCGTGGAGTTTTACCG,

CD34 Fw: GGTAGCTCTCTGCCTGATGAG, CD34 Rev: TGGTAGGAACTGATGGGGATATT,

GATA2 Fw: CACCCCGCCGTATTGAATG, GATA2 Rev: CCTGCGAGTCGAGATGGTTG,

Gsk3b Fw: CTTTGGAAGTGCAAAGCAG, Gsk3b Rev: CCAACTGATCCACACCAC,

Mfsd2a Fw: CTCCTGGCCATCATGCTCTC, Mfsd2a Rev: GGCCACCAAGATGAGAAA,

ZO-1 Fw: GCCGCTAAGAGCACAGCAA, ZO-1 Rev: TCCCCACTCTGAAAATGAGGA,

VE-Cadherin Fw: AACCATGACAACACCGCCA, VE-Cadherin Rev: CGTTGTCTGAGATGAGCACG,

CAV1 Fw: GCGACCCCAAGCATCTCAA, CAV1 Rev: ATGCCGTCGAAACTGTGTGT,

CD93 Fw: GATGGCTCTTTCTACTGCTCCTG, CD93 Rev: CCACACCTGAAGGAACCATCTG,

PTEN Fw: TGAGTTCCCTCAGCCATTGCCT, PTEN Rev: GAGGTTTCCTCTGGTCCTGGTA,

ABCG2 Fw: CAGTTCTCAGCAGCTCTTCGAC, ABCG2 Rev: TCCTCCAGAGATGCCACGGATA,

Tcf4 Fw: CACTTTCCCTAGCTCCTTCTTC, Tcf4 Rev: GTTCGTGTGGTCAGGAGAATAG,

Ck1a1 Fw: TAGCTGACCAGATGATCAG, Ck1a1 Rev: GTATCGGGCAGTGCCAGTG
